# Sex-biased transcriptional rewiring of neuronal circuits associated with eprinomectin resistance in *Haemonchus contortus*

**DOI:** 10.64898/2026.07.20.739568

**Authors:** Rémy Bétous, Robin Lioutaud, Julie Petermann, Sophie Jouffroy, Philippe Jacquiet, Eva Guchen, Elise Courtot, Lea Bordes, Fabrice Guegnard, Guillaume Sallé, Stephen R. Doyle, Anne Lespine

## Abstract

Anthelmintic resistance poses a critical threat to livestock health, with clinical failures of eprinomectin and other macrocyclic lactones documented in the gastrointestinal parasite *Haemonchus contortus* worldwide. We performed whole-genome Pool-seq of larvae and RNA-seq on adult male and female worms from five phenotypically characterised isolates – two eprinomectin-susceptible and three resistant – from dairy sheep in Southwest France. We identified a conserved macrocyclic lactone resistance locus on chromosome 5 and a putatively eprinomectin-specific locus on chromosome 4, alongside sex-specific transcriptomic responses linking genomic selection signatures to transcriptional phenotype. Predicted functions of selected and differentially expressed genes support reorganisation of neuronal circuits in drug responsiveness, including the kynurenine pathway producing neuroactive metabolites, further supported by quinolinic acid potentiating eprinomectin efficacy in larval assays. Despite geographic proximity, resistance appears to have evolved independently across farms. Together, these findings reveal a complex genetic architecture underlying eprinomectin resistance with implications for targeted diagnostics and resistance management.

## Introduction

Parasitic nematodes significantly impact animal health and cause substantial economic losses in the livestock industry^1^. Among these, the gastrointestinal nematode *Haemonchus contortus* is a pervasive parasite of small ruminants across temperate and tropical regions worldwide. Infections pose a serious threat due to the parasite’s high pathogenicity, prolific reproduction, and high genetic diversity^2,3^.

Furthermore, *H. contortus* is a thermophilic species, and climate change is facilitating its spread into new areas. In the Southwest of France, for instance, *H. contortus* infections have recently spread to small ruminants grazing in the mountainous pastures of the Pyrenees, disrupting sheep cheese and meat production at higher altitudes^4,5^.

The control of parasitic nematodes relies almost exclusively on anthelmintic drugs. However, the widespread and rapid emergence of drug resistance has become a critical global threat, undermining the foundation of effective parasite control. For decades, benzimidazoles were the most important anthelmintic class used in livestock, until resistance rendered them ineffective. In response, macrocyclic lactones – avermectins (e.g. ivermectin, eprinomectin) and milbemycins (e.g. moxidectin) – gradually replaced benzimidazoles and are now the most widely used compounds. Among them, eprinomectin is particularly important as the last effective anthelmintic approved for use in dairy animals with a zero-withdrawal period, ensuring that animal-derived products remain safe for immediate human consumption. However, resistance to eprinomectin has emerged in the Southwest of France, a region pivotal to dairy sheep farming and the production of Protected Designation of Origin (PDO) cheeses. The accelerating resistance of *H. contortus* to anthelmintics in this region poses a serious risk to both animal health and the regional agricultural economy^5–8^, mirroring the spread of anthelmintic resistance across Europe and other regions of the world^1–3^.

Identifying and mitigating anthelmintic resistance in parasites like *H. contortus* is one of the most pressing challenges facing the small ruminant livestock industry, yet the genetic mechanisms underlying its evolution remain incompletely understood. The best-characterised resistance variants are point mutations in β-tubulin isotype 1 at codons 167, 198, and 200, which reduce benzimidazole binding affinity^9–11^. For macrocyclic lactones, resistance mechanisms are less clearly defined and appear considerably more complex. Although ivermectin targets glutamate-gated chloride channels (GluCl), and resistance-associated GluCl variants are well-documented in the free-living nematode *Caenorhabditis elegans*^12–15^, evidence for analogous selection at these channels in parasitic nematodes is limited. ABC transporters and amphid development genes have been proposed as candidates^16,17^, though whether they are under direct selection in parasitic species in the field remains unclear.

Genome-wide approaches have begun to refine the signatures of drug-mediated selection^18^; for example, a single major quantitative trait locus (QTL) has been identified on *H. contortus* chromosome 5 consistently associated with ivermectin resistance across controlled genetic crosses^19–22^ and field isolates^23,24^. This region also shows evidence of selection in moxidectin-treated populations, though with recessive rather than dominant inheritance^19^, suggesting that even drugs within the same class may operate through distinct genetic mechanisms. At the transcriptomic level, ivermectin exposure induces broad-scale changes in neuronal plasticity and chloride homeostasis^20^, and changes in expression of *pgp* and cytochrome genes – whose products are expected to reduce drug concentration at the target site – have been a major focus of resistance research^25^, with a *pgp-9* copy number variant associated with ivermectin resistance in *Teladorsagia circumcincta*^26,27^. For eprinomectin – which is the preferred treatment for dairy sheep and has distinct pharmacokinetic properties relative to other macrocyclic lactones – the specific genomic basis of resistance remains largely uncharacterised.

Here, we characterise the genetic and transcriptomic basis of eprinomectin resistance in a geographically localised natural host-parasite system. Using phenotypically characterised *H. contortus* isolates from five dairy sheep farms in the Pyrénées-Atlantiques of southwest France (Fig. 1), we applied whole-genome Pool-seq on larvae and RNA-seq on adult male and female worms. These data identified a conserved macrocyclic lactone resistance locus on chromosome 5 and a putatively eprinomectin-specific locus on chromosome 4, alongside sex-specific transcriptomic responses linking genomic selection signatures to transcriptional phenotype. Functional assays further implicated the kynurenine pathway as a candidate determinant of eprinomectin susceptibility, revealing a polygenic resistance architecture with distinct sex-dependent transcriptional consequences.

**Fig. 1:**
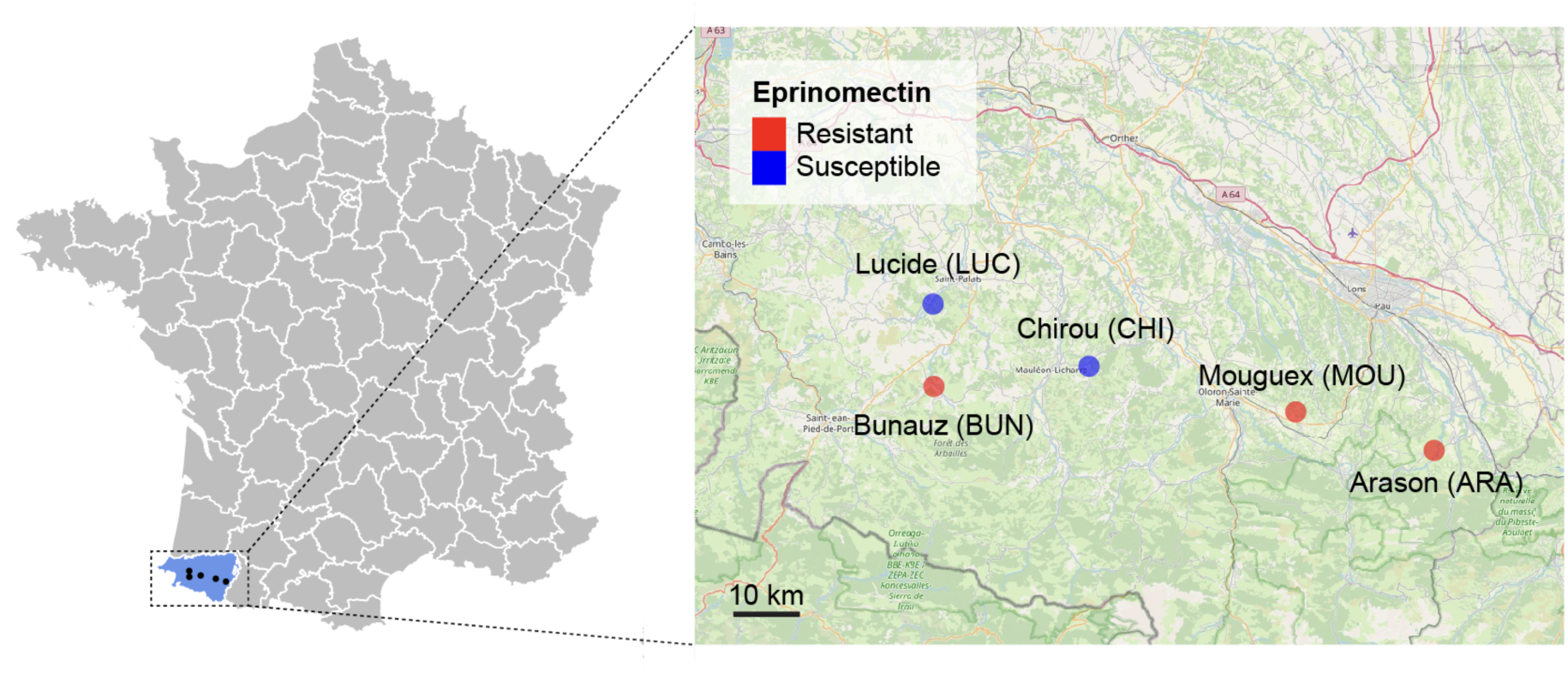
Geographic location of farms where eprinomectin-susceptible and resistant *Haemonchus contortus* populations were collected. *H. contortus* isolates were collected from five farms located within a geographically restricted area of Southwest France (∼3,000 km²). Two farms were classified as eprinomectin-susceptible (Lucide, LUC; Chirou, CHI; blue) and three as resistant (Bunauz, BUN; Mouguex, MOU; Arason, ARA; red). Classification was based on prior assessment of field efficacy using faecal egg count reduction tests (FECRT)^4,5^ and larval motility measured with the Worm MicroTracker^7^.

## Results

### Phenotypic characterisation of *Haemonchus contortus* isolates

We selected five dairy sheep farms (ARA, BUN, CHI, LUC, and MOU) in southwest France (Fig. 1) based on their routine use of eprinomectin as the primary anthelmintic. Previous field assessments confirmed eprinomectin therapeutic failure on three farms (ARA, BUN, MOU), where faecal egg count reduction tests (FECRT — a standard measure of anthelmintic efficacy based on the proportional reduction in strongylid egg output post-treatment) fell below the 95% efficacy threshold despite plasma drug concentrations consistent with adequate exposure^5,7^. In contrast, FECRT values reached 100% at CHI and LUC, indicating full clinical efficacy. *In vitro* larval motility assays on infective larvae (L3) confirmed CHI and LUC as fully susceptible to eprinomectin, ivermectin, and moxidectin, while ARA, BUN, and MOU were resistant to all three, with eprinomectin resistance factors (RFs) ranging from 51 to 109 – substantially higher than those for ivermectin and moxidectin (Supplementary Table 1). These populations were accordingly classified as macrocyclic lactone-resistant^7^. Thiabendazole assessment revealed benzimidazole resistance in ARA and MOU; BUN showed no evidence of benzimidazole resistance (RF = 0.5), and CHI and LUC remained fully susceptible (Supplementary Table 1).

### Candidate genomic regions of drug-mediated selection between susceptible and eprinomectin-resistant isolates

To assess the genetic impact of eprinomectin treatment, we performed whole-genome Pool-seq on pooled L3 larvae from each isolate (Supplementary Data 1), generating 638 million reads at a mean coverage of 53.3× per isolate (Supplementary Data 2). Variant calling identified an average of 6.5 million SNPs and 571,000 indels per sample, of which 94.2% were shared between at least two isolates (Supplementary Data 2). Pairwise genome-wide *F*_ST_ comparisons showed that CHI and LUC were genetically most similar (*F*_ST_ = 0.008), while ARA was the most genetically distinct among resistant isolates, consistent with its more easterly geographic position (Supplementary Fig. 1). Sliding-window *F*_ST_ scans comparing susceptible and resistant populations identified five candidate QTLs of eprinomectin-mediated selection: one peak each on chromosomes 1, 2, and 4, and two peaks on chromosome 5 (Fig. 2a).

**Fig. 2:**
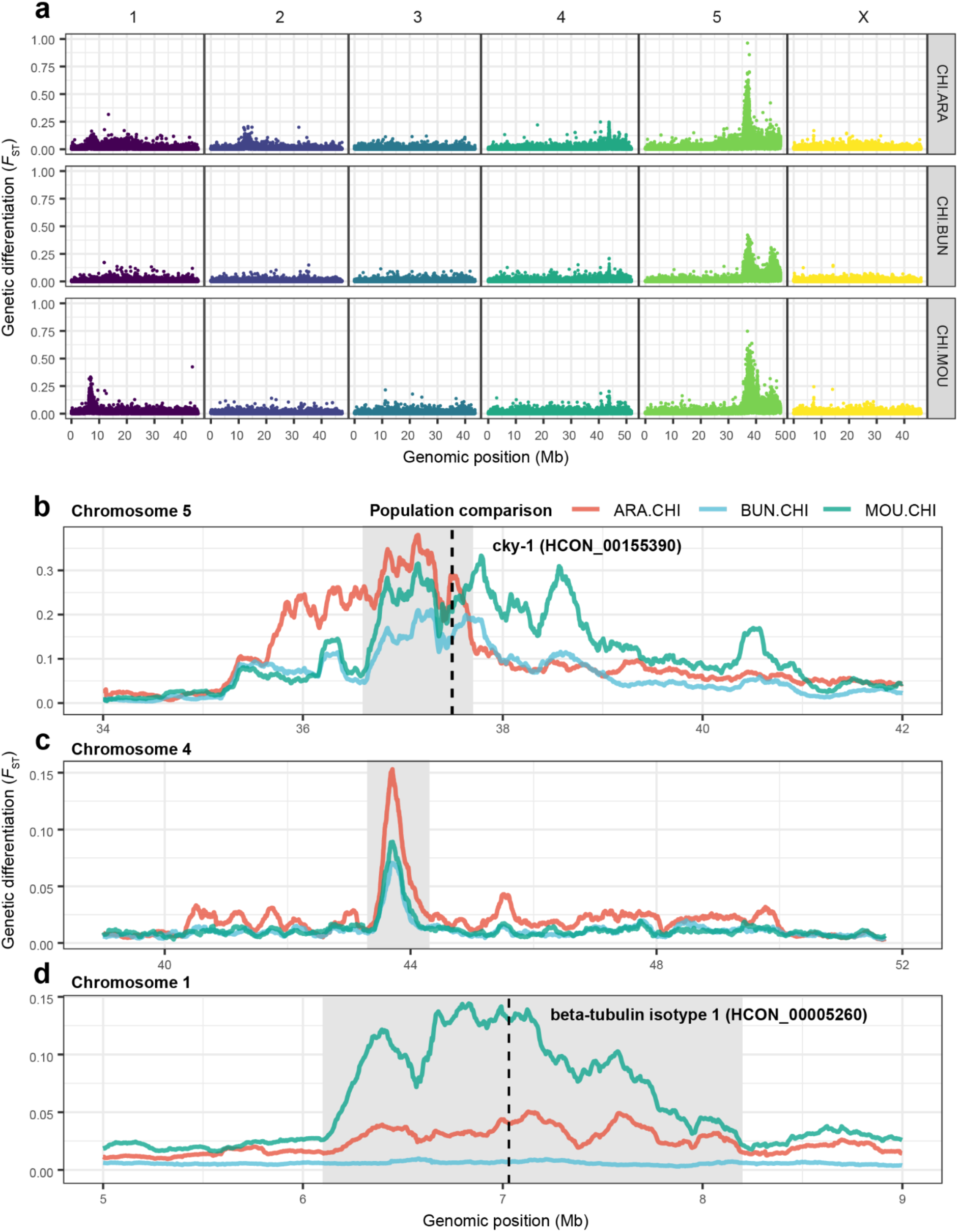
Genome-wide analysis of genetic differentiation between eprinomectin-susceptible and resistant *Haemonchus contortus* populations. Pairwise *F*ST between eprinomectin-susceptible (CHI) and resistant (ARA, BUN, MOU) isolates. **a** Genome-wide Manhattan plots showing pairwise *F*ST between the susceptible CHI isolate and each resistant isolate. Each point represents the mean *F*ST in non-overlapping 5,000-bp windows with 2,500-bp overlap. **b–d** Zoomed views of selected peaks, showing smoothed *F*ST (200-kb sliding windows, computed from 5,000-bp windows with 2,500-bp overlap) for pairwise comparisons between each resistant isolate and the susceptible CHI isolate. Colours represent distinct pairwise comparisons (legend, bottom). Grey-shaded regions indicate the consensus peak boundaries across comparisons. **b** Major differentiation peak on chromosome 5, encompassing *cky-1* (HCON_00155390), a gene within the QTL region previously associated with ivermectin resistance. **c** Peak on chromosome 4, driven primarily by the BUN isolate, putatively eprinomectin-specific. **d** Peak on chromosome 1 centred on *β-tubulin isotype 1* (HCON_00005260), a well-characterised benzimidazole resistance locus; consistent with phenotypic data, this peak is absent in the BUN comparison, which showed no benzimidazole resistance.

A dominant *F*_ST_ peak shared across all pairwise comparisons was identified on chromosome 5 (consensus region ∼36.6–37.7 Mb), coinciding with a locus repeatedly identified in ivermectin resistance studies^19,21,22^. Filtering for variants at low frequency in susceptible isolates (AF < 0.1) that rose to high frequency in resistant isolates (AF > 0.4) resolved a narrow peak spanning ∼36.73–37.5 Mb (Fig. 3a), with 38 genes showing mean ΔAF > 0.7 between resistant and susceptible isolates (Supplementary Data 3). Twenty-five of these genes harbour at least one variant predicted to have high or moderate functional impact (SnpEff), of which nine have putative *C. elegans* orthologs.

**Fig. 3:**
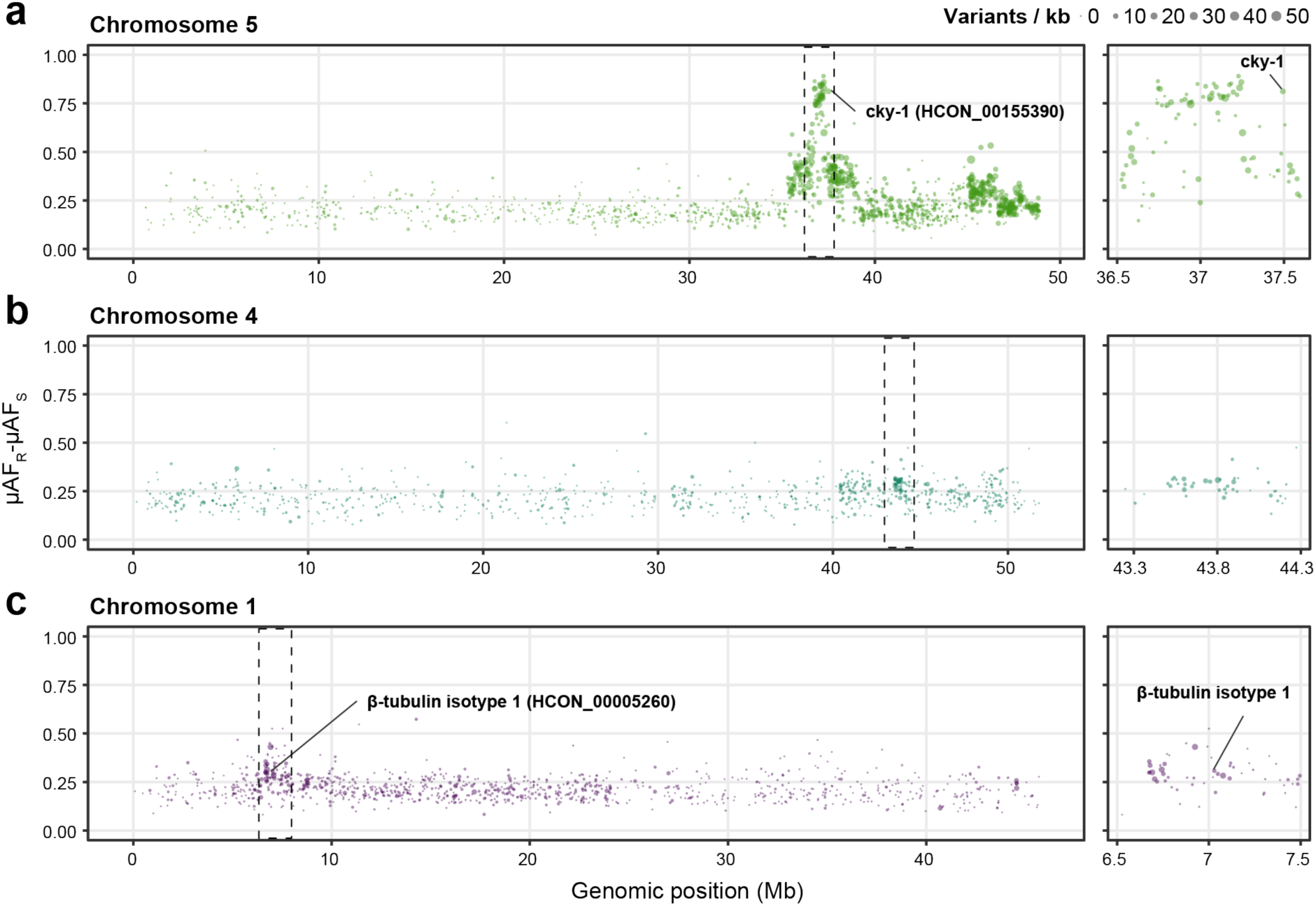
Gene-wise difference in mean allele frequency between susceptible and eprinomectin-resistant isolates on chromosomes 5, 4, and 1. The y-axis shows ΔAF = μAFR − μAFS, the difference in mean allele frequency between resistant and susceptible isolates. Each point represents a gene containing at least one qualifying variant, positioned at its midpoint along the chromosome (x-axis). Point size reflects the number of variants per gene per kilobase. Dashed boxes indicate the peak regions shown in the zoomed insets (right). a Chromosome 5, with *cky-1* (HCON_00155390) annotated as the top candidate. b Chromosome 4. c Chromosome 1, with the position of *β-tubulin isotype 1* (HCON_00005260), a well-characterised benzimidazole resistance locus, annotated.

The strongest candidate is *cky-1* (HCON_00155390; ΔAF = 0.84), a transcription factor whose increased expression has been associated with ivermectin resistance in *H. contortus*^20,21^. Other prominent candidates include *sulp-8* (*HCON_00155080*; ΔAF = 0.82), predicted to encode a chloride/sulfate transmembrane transporter, and *ido* (HCON_00155060; ΔAF = 0.78), encoding an indoleamine 2,3-dioxygenase that initiates the kynurenine pathway. Two GPCRs with moderate-impact variants were also identified: HCON_00154994 (ΔAF = 0.80) and HCON_00154992 (ΔAF = 0.78), the latter orthologous to *C. elegans str-257*.

Towards the end of chromosome 5, a second minor peak of genetic differentiation is visible (Fig. 2a), particularly prominent in the CHI–BUN comparison. This region has been previously identified as differentiated in backcross-derived ivermectin-resistant isolates^22^; however, neither prior analyses nor our data resolve a discrete region of differentiation clearly distinct from the main chromosome 5 QTL. The region contains orthologs of *avr-15* (HCON_00161180; 45.13 Mb), encoding a glutamate-gated chloride channel subunit directly implicated in macrocyclic lactone mode of action, and *pgp-11* (HCON_00162780; 47.24 Mb), encoding a P-glycoprotein previously associated with ivermectin resistance in *C. elegans* and other nematode parasites.

However, no significant differences in variant frequency were detected in this region between resistant and susceptible isolates, suggesting these genes are not under strong selection and may play at most a minor role in eprinomectin resistance.

A peak of genetic differentiation is also observed on chromosome 4 (Figures 2a), spanning ∼43.3–44.3 Mb (Figures 2c), with the strongest signal in the ARA isolate. This signature of selection was not observed in previous genomic studies of ivermectin- or moxidectin-resistant *H. contortus*, suggesting it may reflect eprinomectin-specific selective pressure. The region contains 99 genes, including candidates potentially involved in chemoreception (8 GPCRs) and drug detoxification (6 UDP-glucuronosyltransferases, UGTs phase II metabolic enzymes that could contribute to drug inactivation) (Supplementary Data 4). Within a narrower subregion (∼43.5–43.9 Mb), we identified a local enrichment of high-frequency variants (Fig. 3b), containing five genes with more than five high-frequency variants per kilobase. Two of these have *C. elegans* orthologs (HCON_00125700/*C52G5.2* and HCON_00125765/*R09H10.6*), though their functions remain uncharacterised. The accumulation of high-frequency variants within this region is consistent with eprinomectin-mediated selection, though contributions from other evolutionary or demographic processes cannot be excluded.

Peaks of genetic differentiation centred on *β-tubulin isotype 1* (HCON_00005260) were observed on chromosome 1 in two resistant isolates, ARA and MOU (Figures 2a,d), consistent with their benzimidazole-resistant phenotypes (Table 1, Supplementary Data 1). The *F*_ST_ peak surrounding *β-tubulin isotype 1* was greater in MOU than ARA, consistent with a greater allele frequency differentiation at this locus in MOU (ΔAF: MOU = 0.66, ARA = 0.34) relative to the anthelmintic-susceptible CHI and LUC and BUN isolates (AF ≈ 0.03–0.04; Fig. 3c). Genotyping of known resistance-associated variants revealed Glu198Ala and Phe200Tyr at moderate frequencies in ARA (35.4% and 41.5%, respectively), whereas in MOU, Glu198Ala predominated (75.6%) over Phe200Tyr (14.0%; Supplementary Data 5). The higher benzimidazole resistance observed in MOU (Supplementary Data 1) is consistent with prior evidence that Glu198Ala confers greater resistance than Phe200Tyr^28^.

Together, these patterns likely reflect farm-level differences in historical benzimidazole use prior to the introduction of eprinomectin in 2016. Notably, Phe200Tyr in *β-tubulin isotype 1* was also detected at low frequency in BUN, CHI, and LUC (13.8%, 6.2%, and 7.1%, respectively), suggesting these populations retain the genetic potential to develop benzimidazole resistance under renewed selection pressure.

In the ARA isolate, an additional peak was observed on chromosome 2 (Fig. 2a), coinciding with β-tubulin isotype 2 (HCON_00043670; Supplementary Fig. 2).

Variants in this gene have been associated with high-level benzimidazole resistance when co-occurring with isotype 1 variants^11,21^. However, the previously reported Glu198Val variant in isotype 2 was absent; instead, we identified a His6Leu substitution in both ARA and MOU at frequencies of 19.1% and 22.7%, respectively. Structurally, this substitution lies approximately three β-sheets from the benzimidazole binding site and is not an obvious causal variant, though its contribution to resistance remains undetermined.

Taken together, these findings confirm that benzimidazole susceptibility persists in some isolates and that β-tubulin variants were selected under historical farm-level drug pressure and maintained over time.

### Eprinomectin resistance has likely evolved independently on geographically close farms

The close geographic proximity of susceptible and resistant populations prompted us to ask whether eprinomectin resistance arose once and spread among farms, or evolved independently. Genome-wide comparisons between susceptible populations revealed very low genetic differentiation (Supplementary Fig. 3), suggesting a shared ancestral background. Pairwise comparisons among resistant populations revealed elevated differentiation at the same loci identified in susceptible-vs-resistant comparisons (Supplementary Fig. 4), consistent with independent selection of resistance haplotypes in each population; had resistance spread from a single origin, resistant populations would be expected to share haplotypes at these loci and show reduced inter-resistant differentiation. The ARA population was the most differentiated, consistent with its distinct genomic background at other loci genome-wide. Benzimidazole resistance genetics further support independent selection histories across farms, evidenced by different β-tubulin variants in ARA and MOU (Supplementary Data 5). Notably, the chromosome 5 eprinomectin-associated peak showed greater differentiation between ARA and either MOU or BUN than between BUN and MOU, suggesting BUN and MOU may share a common resistance haplotype at this locus. The chromosome 4 peak was not detectable in any inter-resistant comparison. Together, these data suggest that resistance to eprinomectin and benzimidazoles has evolved independently on at least two occasions across these farms, through partly distinct genetic routes.

### Drug-resistant isolates have common and sex-specific differential gene expression that is enriched in selected QTLs

Transcriptional differences between susceptible and eprinomectin-resistant isolates were assessed using RNA-seq from adult male and female worms (Supplementary Data 1). On average, 54.8 million reads were obtained per sample, with approximately 78.8% and 81.2% mapping to the *H. contortus* reference genome for males and females, respectively (Supplementary Data 6). Principal component analysis showed samples clustered by sex along PC1 (62.29% of variance), with susceptible and resistant isolates separating along PC2 (3.41%; Fig. 4a), reflecting the dominant effect of biological sex on global transcription. Sex-stratified PCAs confirmed transcriptomic separation between susceptible and resistant isolates in both sexes independently (Supplementary Fig. 5).

**Fig. 4:**
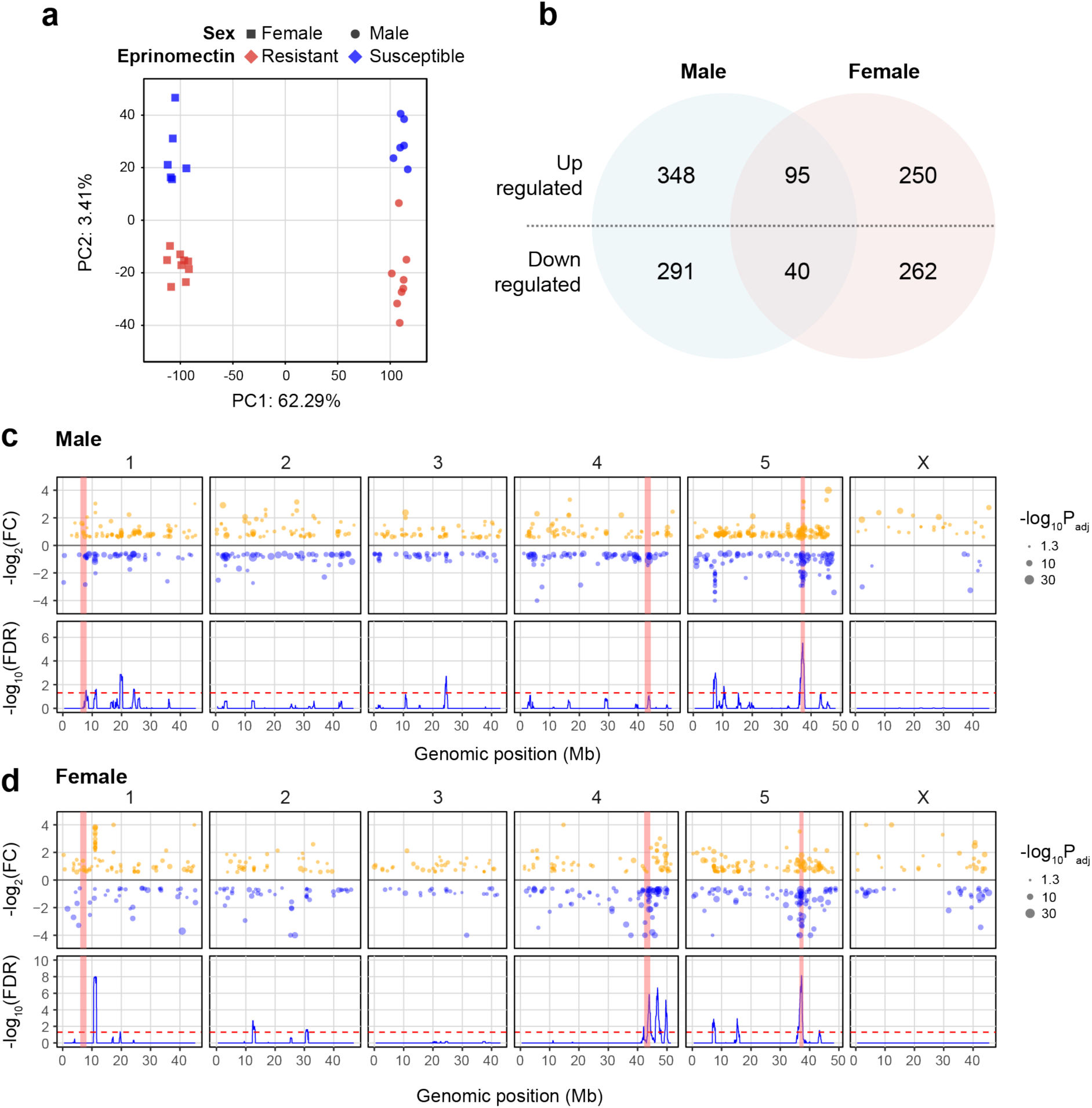
Transcriptomic comparison of susceptible and drug-resistant isolates reveals shared and sex-specific differential expression enriched within selection QTLs. a Principal component analysis (PCA) based on the expression of 14,112 genes separates samples by sex along PC1 (62.29% of variance) and by resistance status along PC2 (3.41%), reflecting the dominant effect of biological sex on global gene expression and motivating sex-stratified analysis. b Venn diagram showing differentially expressed genes (DEGs) between susceptible and resistant isolates in males, females, and both sexes combined. c, d Genome-wide karyoplots of DEGs in c resistant males and d resistant females. Upper tracks: yellow and blue points indicate genes significantly upregulated or downregulated in resistant relative to susceptible isolates (adjusted p < 0.05); point size reflects statistical significance; fold-change values outside [−4, 4] were truncated to the boundary values for visualisation. Lower tracks: blue lines show −log₁₀(FDR) from hypergeometric enrichment tests assessing clustering of DEGs within 1 Mb sliding windows (100 kb step); the red dashed line indicates FDR = 0.05 (Benjamini–Hochberg correction). Red shaded boxes indicate the positions of genomic differentiation peaks from Fig. 2, shown at identical positions in both panels for reference.

Differential expression analysis between susceptible and resistant isolates identified 2,851 and 2,508 DEGs in males and females, respectively (adjusted p < 0.05). After applying an |FC| ≥ 1.5 threshold, these are reduced to 774 DEGs in males and 647 in females (Fig. 4b, Supplementary Data 7). The large majority were sex-specific (639 in males, 512 in females), with 135 shared between sexes (95 upregulated, 40 downregulated).

We next examined the genomic distribution of DEGs to assess whether their localisation deviated from random expectations across the five autosomes and the X chromosome (Fig. 4c, d). Hypergeometric testing confirmed significant non-random genome-wide distribution; DEG-enriched regions on chromosome 5 overlapped the major genomic QTL (∼36.6–37.7 Mb) in both males (∼35.5–38.3 Mb; n = 45 DEGs) and females (∼35.1–38.2 Mb; n = 43 DEGs), and together with a strong, female-specific enrichment of 73 DEGs across chromosome 4 (∼41.5–50.5 Mb), surrounding the eprinomectin-associated differentiation peak. A smaller enriched region on chromosome 1 in females (10.1–12.0 Mb; 17 upregulated, 2 downregulated) contained 14 *sip-1* homologs, suggesting a transcriptional stress response independent of the benzimidazole selection signal. In total, 141 DEGs (∼11% of all DEGs) fell within genomic selection QTLs.

### Conserved and eprinomectin-specific transcriptomic signatures across macrocyclic lactones

To determine whether the observed transcriptomic signatures were specific to eprinomectin resistance, we applied the same analysis pipeline to published RNA-seq data from experimentally selected ivermectin- and moxidectin-resistant *H. contortus* lines^19^. Karyoplot analysis revealed significant DEG enrichment at ∼34.9–46.7 Mb on chromosome 5 in both ivermectin- and moxidectin-resistant worms of both sexes, overlapping the chromosome 5 QTL — a broader region than in eprinomectin-resistant isolates, likely reflecting differences in selection history between experimentally selected lines and field-derived populations. In contrast, no DEG enrichment was detected at the chromosome 4 locus in either drug or sex (Supplementary Fig. 6), confirming that the chromosome 4 signal is female-specific and unique to eprinomectin resistance.

The colocalisation of the chromosome 5 QTL and DEG enrichment prompted examination of *cky-1*, a transcription factor previously implicated in ivermectin resistance^20,21^ via regulation of neuronal excitability. No reads mapping to *cky-1* were detected in RNA-seq data from any sample, likely reflecting low expression of a tightly regulated transcription factor. However, RT-qPCR confirmed *cky-1* expression in all sample sets, with a modest but statistically significant reduction of expression in resistant females and males (Supplementary Fig. 7); this is in contrast to the previous finding of significant overexpression of *cky-1* in ivermectin-resistant parasites^20,21^. Proposed downstream targets of *cky-1* were significantly downregulated, including *pmk-1* and *pmk-2* (p38 MAP kinases; both sexes) and *vab-3* (Pax transcription factor; females only), suggesting residual *cky-1* regulatory activity despite low expression levels. These findings suggest that *cky-1*-mediated transcriptional dysregulation may contribute to macrocyclic lactone resistance, potentially through modulation of stress signalling and neuronal development pathways.

### Transcriptomic signatures of resistance implicate neuronal, transport, and structural pathways

To understand the global effects of eprinomectin resistance on gene expression, we performed GO enrichment analysis on male and female DEG sets independently before identifying terms common to both sexes (Fig. 5a, b; Supplementary Data 8). In both sexes, "extracellular region" (GO:0005576) was the most significantly enriched term (108 DEGs in males, 46 in females), the majority being orthologs of *C. elegans vap-1*, a cysteine-rich secretory protein involved in extracellular matrix remodelling around sensory neurons implicated in macrocyclic lactone uptake^29^. In *C. elegans*, *vap-1* expression is regulated by the transcription factor PROS-1^29^, whose two putative *H. contortus* orthologs (HCON_00075020 and HCON_00075030) were both significantly upregulated in eprinomectin-resistant males. Peptidase and endopeptidase activities (GO:0008233, GO:0004175) were also enriched in both sexes, with contributing genes encoding secreted peptidases previously identified in extracellular vesicles and implicated in host-pathogen communication^30^ (Supplementary Data 8); notably, HCON_00101590 appeared in both sexes, making it a candidate of particular interest.

**Fig. 5:**
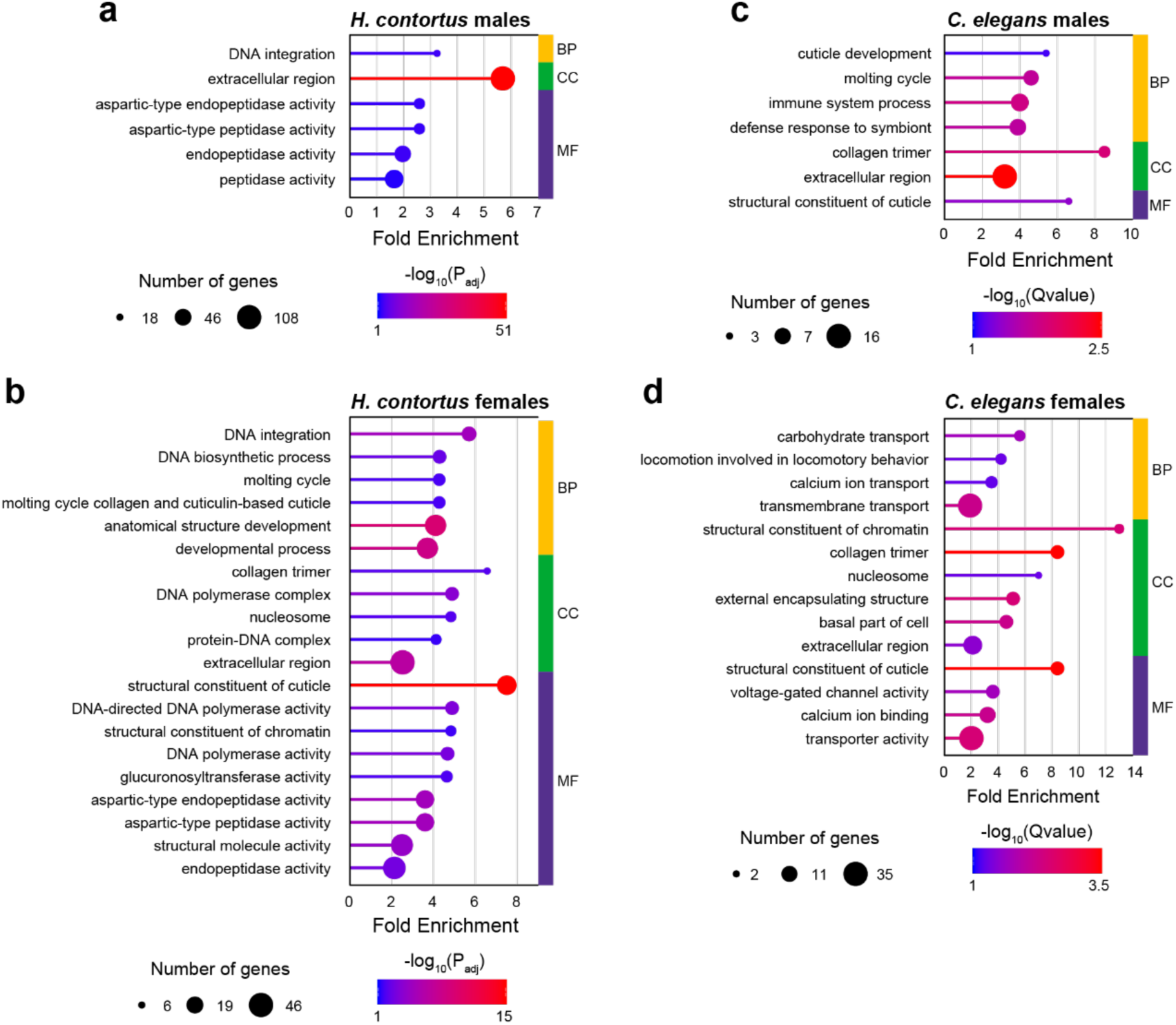
Gene ontology enrichment of differentially expressed genes reveals distinct and shared pathways in male and female eprinomectin-resistant isolates. Significantly enriched Gene Ontology (GO) terms from differentially expressed genes between eprinomectin-susceptible and eprinomectin-resistant *H. contortus* isolates. a, b GO terms enriched in *H. contortus* males and females, respectively. c, d GO terms enriched among *C. elegans* orthologs of differentially expressed genes in males and females, respectively. The x-axis shows fold enrichment; dot size reflects the number of genes contributing to each GO term; dot colour indicates statistical significance (−log10(adjusted p-value) for a, b; −log10(Q-value) for c, d), with warmer colours indicating greater significance. Coloured bars on the right of each panel indicate the GO category: BP, biological process; CC, cellular component; MF, molecular function.

Using reciprocal best-hit ortholog mapping to *C. elegans* (9,025 pairs), GO overrepresentation analysis identified cuticle structure (GO:0042302) as enriched in both sexes, and transmembrane transport (GO:0055085) prominently in females (Fig. 5c, d; Supplementary Data 9). Within the transmembrane transport category, *pgp-9.1* and *pgp-9.2* – putative macrocyclic lactone efflux transporters – were strongly upregulated in resistant females, together with *pgp-2* (HCON_00004450) and *pgp-3* (HCON_00042800), while *pgp-1* (HCON_00098130) was downregulated; *pgp-9.1* was also upregulated in resistant males, though to a lesser extent. This pattern is consistent with increased drug efflux in resistant worms.

Voltage-gated ion channels, which regulate action potentials, neurotransmitter release, and synaptic transmission in *C. elegans*^31^, were also represented: *unc-36* (HCON_00118530; voltage-gated calcium channel) was strongly downregulated in both sexes, as were orthologs of *twk-24* and *twk-46* (two-pore domain potassium channels; HCON_00163840 and HCON_00161600, respectively). Additionally, HCON_00162485, orthologous to the *sleepless* gene and predicted to regulate voltage-gated potassium channel activity and cholinergic synaptic transmission, was differentially expressed in resistant females, which is notable given the role of macrocyclic lactones in inducing paralysis through neuronal pathways.

Several neurotransmitter-related genes were also differentially expressed. *cho-1* (HCON_00113930) and *glt-1* (HCON_00058990), transporters involved in acetylcholine and glutamate synaptic homeostasis respectively, were strongly upregulated in resistant females. The GABA receptor *gab-1* (HCON_00076450) was strongly downregulated in both sexes, while the acetylcholine receptor *acr-9* (HCON_00188970) was significantly upregulated in females only. The glutamate receptor *glc-5* (HCON_00028600), orthologous to the macrocyclic lactone target HcGluClα^32^, was modestly upregulated in resistant males (FC = 1.33); while unlikely to directly reduce susceptibility, this may reflect broader reorganisation of synaptic receptor expression. Finally, *unc-7* (HCON_00190630), encoding an innexin gap junction subunit, was significantly downregulated in resistant females, consistent with prior evidence that *unc-7* mutations confer ivermectin resistance^33^ and suggesting a female-specific contribution of gap junction signalling to the resistance phenotype.

WormBase phenotype enrichment analysis identified genes linked to "anthelmintic-response variant” phenotype (WBPhenotype:0001852) in resistant males, including *dyf-2* (HCON_00089540) and *osm-12* (HCON_00041970), whose disruption reduces macrocyclic lactone uptake through effects on amphid neuron structure^33^ (Supplementary Data 10). Consistent with this, *osm-12* was among the most strongly downregulated genes in resistant males (FC = −3.3). Among the 16 phenotypes enriched in both sexes, several related to neuronal function, including "chemosensory behaviour" (WBPhenotype:0001049), "neuropil development" (WBPhenotype:0000945), and "ciliated neuron morphology" (WBPhenotype:0001526), the last of which has been repeatedly associated with macrocyclic lactone resistance^16,33^. Eight further enriched phenotypes related to locomotion, including "foraging hyperactive", "forward locomotion increased", and "turning variant", consistent with the expected behavioural consequences of altered neuronal circuitry.

Taken together, these findings point to a broad reorganisation of neuronal circuit function in eprinomectin-resistant worms, encompassing sensory neuron morphology, synaptic transmission, and locomotor behaviour, reflecting changes that likely influence both drug uptake and the downstream neuronal response to macrocyclic lactone exposure.

### Distinct and shared transcriptomic responses between eprinomectin, ivermectin, and moxidectin-resistant isolates

To assess the generalisability of our transcriptomic findings, we reanalysed RNA-seq data from ivermectin- and moxidectin-resistant *H. contortus* lines derived from experimental evolution^19^, identifying DEGs shared across or specific to each drug (Supplementary Fig. 8). The majority of DEGs identified in eprinomectin-resistant isolates did not overlap with those from experimentally evolved lines, suggesting that field-derived resistance generates transcriptional responses that are largely drug-and context-specific. Nevertheless, shared DEGs across all three macrocyclic lactones were enriched for genes in the chromosome 5 QTL region and included candidates involved in drug transport and neuronal function, consistent with a conserved core resistance response.

When upregulated and downregulated genes were considered separately, a clear drug-specific transcriptional signature emerged. Among upregulated genes, eprinomectin-resistant worms showed the strongest and most distinct signature, with 91 DEGs unique to the eprinomectin-resistant field isolates. Moxidectin-resistant worms showed 22 moxidectin-specific upregulated DEGs, whereas no gene was uniquely upregulated in ivermectin-selected worms of either sex. Among downregulated genes, moxidectin-resistant worms showed the most pronounced specificity (50 unique DEGs), followed by eprinomectin-resistant worms (27 unique DEGs), while ivermectin-selected worms again showed minimal specificity (1 unique DEG). The transcriptional signatures in ivermectin-selected crosses likely reflect both lower variation in selection intensity and the constrained genetic background of experimental crosses, which together minimise non-specific transcriptional responses compared to field-derived isolates.

Surprisingly, no gene was upregulated in all three drugs across both sexes. In contrast, three genes – *ido* (HCON_00155060), *sulp-8* (HCON_00155080), and HCON_00155240 – were consistently downregulated across all three resistance backgrounds in both sexes, strongly implicating them in a shared core mechanism of macrocyclic lactone resistance. All three lie within the chromosome 5 genomic QTL and the DEG-enriched region identified by karyoplot analysis (Fig. 5), and two — *ido* and *sulp-8* — showed the highest allele frequency shifts in the genomic analysis (ΔAF = 0.711 and 0.675, respectively), with HCON_00155240 also elevated (ΔAF = 0.428).

HCON_00155240, the only gene previously identified as differentially expressed in ivermectin-resistant *H. contortus*^20^, contains a checkpoint kinase-like domain and may have up to 25 paralogs clustering within the chromosome 5 QTL; the absence of functional characterisation of this gene family currently precludes interpretation of its specific role. *sulp-8* (HCON_00155080), orthologous to *C. elegans sulp-8* and related to the mammalian SLC26 anion transporter family^34^, is a particularly compelling candidate; its consistent downregulation across all three resistance backgrounds, combined with high allele frequency shift, suggests a conserved adaptive response potentially counteracting drug-induced chloride influx at the target site.

### The kynurenine pathway modulates eprinomectin susceptibility in *H. contortus*

The third consistently downregulated gene, HCON_00155060, has no identifiable *C. elegans* ortholog but contains an indoleamine 2,3-dioxygenase (IDO) domain, with IDO1 and IDO2 as its closest mammalian homologs. These enzymes catalyse the first committed step of tryptophan catabolism, converting tryptophan to N-formylkynurenine and initiating the kynurenine pathway (Fig. 6a). Consistent with broader pathway dysregulation, *afmd-1*, encoding the enzyme converting N-formylkynurenine to kynurenine, was also significantly downregulated in resistant females (Fig. 6a, Supplementary Data 11), suggesting coordinated suppression of kynurenine biosynthesis in a sex-dependent manner.

**Fig. 6:**
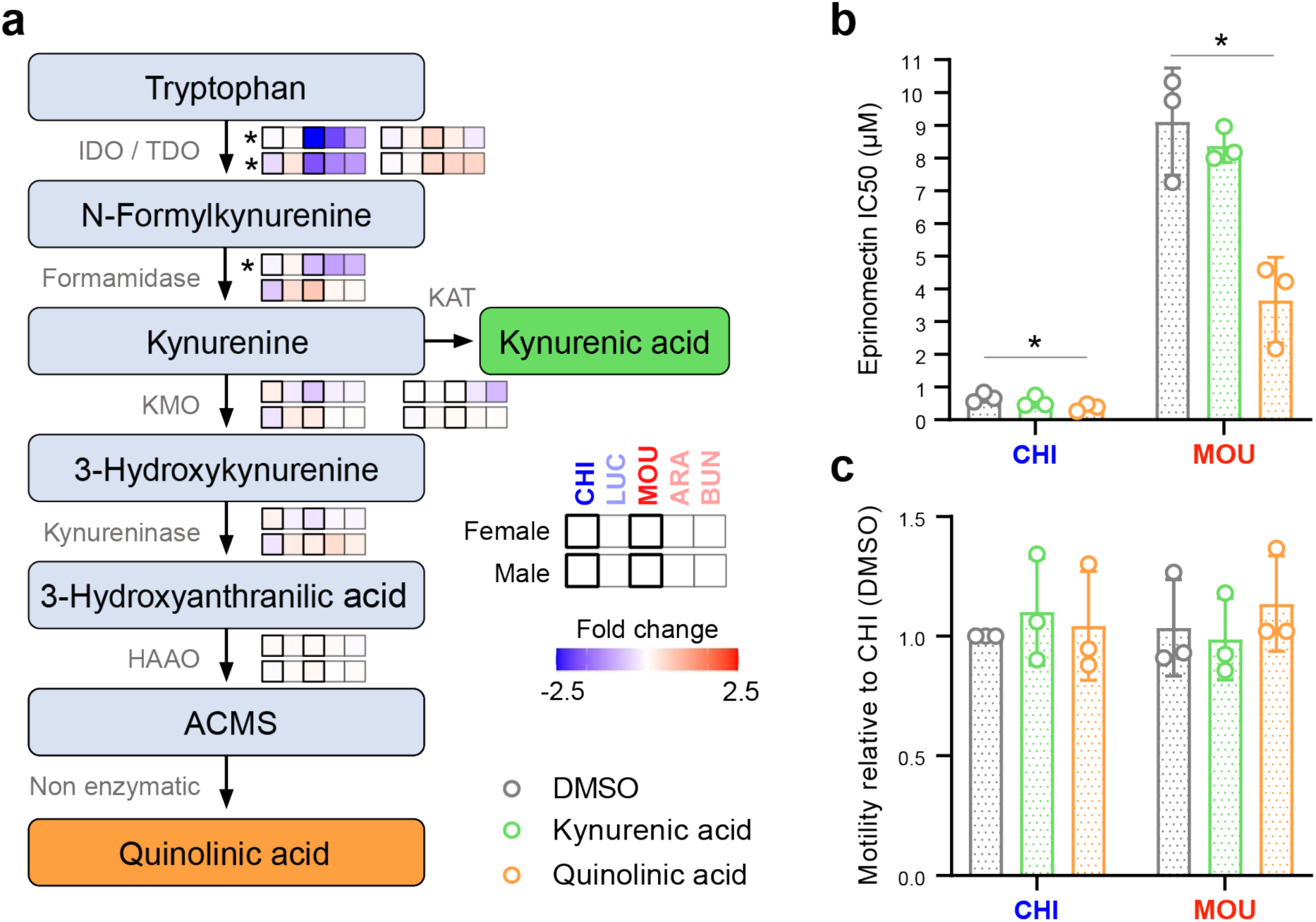
Quinolinic acid modulates eprinomectin susceptibility in *Haemonchus contortus*. **a** Schematic representation of the kynurenine pathway, from tryptophan degradation to quinolinic acid production. Grey text indicates enzyme names or reaction types. Abbreviations: IDO, indoleamine-2,3-dioxygenase; TDO, tryptophan-2,3-dioxygenase; KAT, kynurenine aminotransferase; KMO, kynurenine-3-monooxygenase; HAAO, 3-hydroxyanthranilate 3,4-dioxygenase; ACMS, α-amino-β-carboxymuconate-ε-semialdehyde. Heatmaps show gene expression (fold change in resistant vs susceptible worms, stratified by sex); asterisks on the heatmaps denote differentially expressed genes between susceptible and resistant isolates (adjusted *p* < 0.05). **b** Quinolinic acid potentiates eprinomectin efficacy in resistant larvae. Half-maximal inhibitory concentration (IC₅₀) of eprinomectin following exposure to vehicle (DMSO), kynurenic acid, or quinolinic acid (1 mM each). **c** Kynurenine pathway intermediates do not impair baseline larval motility, ruling out general toxicity. Motility normalised to CHI + DMSO control. Each point represents the mean of technical replicates per experiment; bars show mean ± SD (*n* = 10 wells from 3 independent experiments). \**p* < 0.05 (unpaired *t*-test between DMSO control and subject).

Several kynurenine pathway metabolites possess glutamate receptor-modulating properties relevant to macrocyclic lactone mode of action. Kynurenic acid acts as a competitive antagonist at ionotropic glutamate receptors (AMPA, kainate, and NMDA), preferentially attenuating NMDA receptor activity at the glycine co-agonist site^35^, and modulates neuronal excitability and synaptic plasticity^36^. Quinolinic acid acts as a selective NMDA receptor agonist^37^. Both metabolites influence extracellular glutamate levels and glutamatergic neurotransmission in mammals^38^, functions potentially relevant to *H. contortus*, where macrocyclic lactones act on glutamate-gated chloride channels. We therefore investigated whether these metabolites alter larval eprinomectin susceptibility (Fig. 6b, Supplementary Fig. 9, Supplementary Data 12). Kynurenic acid did not significantly alter the eprinomectin IC₅₀ in either isolate. In contrast, quinolinic acid significantly reduced the IC₅₀ in both susceptible (CHI) and resistant (MOU) larvae, suggesting an effect on the drug target pathway itself rather than specifically on resistance mechanisms. Quinolinic acid alone had no effect on larval motility (Fig. 6c), confirming that sensitisation reflects specific modulation of eprinomectin susceptibility rather than general toxicity. Together, these findings identify the kynurenine pathway as a contributor to eprinomectin susceptibility, with downregulation of *ido* and *afmd-1* in resistant worms potentially reducing quinolinic acid availability and attenuating glutamatergic signalling at the drug target.

## Discussion

Here, we integrate genomic and transcriptomic data to characterise naturally occurring eprinomectin resistance in *H. contortus*. While previous work has characterised ivermectin and moxidectin resistance under experimental or backcross designs^19–22^, our study combines clinical and laboratory phenotyping with whole-genome Pool-seq and sex-stratified RNA-seq on field-derived isolates. The close geographic origin of the resistant isolates minimised background genomic variation, substantially increasing power to detect shared resistance signals. Comparison against published ivermectin- and moxidectin-selected datasets allowed us to distinguish signals shared across macrocyclic lactones from those specific to eprinomectin or local population history. Together, these findings support a model of macrocyclic lactone resistance in which metabolic and neuronal adaptations operate alongside changes at drug target loci.

Genetic differentiation and transcriptomic analyses identified major QTLs on chromosomes 4 and 5 associated with eprinomectin resistance, with significant DEG enrichment within or around both regions, though the two loci differ markedly in their evolutionary context and drug specificity. The chromosome 5 QTL overlaps a locus repeatedly associated with ivermectin and moxidectin resistance^19,20^, supported consistently by genetic differentiation, allele frequency, and DEG enrichment analyses. Cross-species transcriptomic comparison identified only three genes consistently dysregulated across all three macrocyclic lactone resistance backgrounds: *ido*, *sulp-8*, and *HCON_00155240*. Of these, *ido* and *sulp-8* also showed elevated variant allele frequencies in resistant isolates (ΔAF = 0.711 and 0.675, respectively), making them the strongest convergent candidates within this shared locus. All three were consistently downregulated, suggesting that resistance may involve suppression of specific physiological processes rather than upregulation of classical detoxification genes. The chromosome 4 QTL, by contrast, has not been reported in ivermectin- or moxidectin-selected populations, raising the possibility that it represents an eprinomectin-specific selective target. The strong DEG enrichment extending beyond the differentiation peak was predominantly female-specific, consistent with regulatory variants acting across a broader chromosomal domain, though local population structure, linked selection, and sex-specific tissue composition cannot be excluded. Candidate genes within this region include GPCRs, of interest given that ivermectin inhibits several GPCRs at low micromolar concentrations^39^, and UGTs, phase II detoxification enzymes increasingly implicated in anthelmintic response variation^40^. Multiple kynurenine pathway components also map to this region: *aat-1* (*HCON_00125930*), a transporter required for neuronal tryptophan uptake and kynurenic acid synthesis in *C. elegans*^41^, lies within the QTL at 44.2 Mb, while *afmd-1* (*HCON_00129435*), encoding the formamidase immediately downstream of IDO, was significantly downregulated in resistant females and maps nearby at 49.6 Mb. Together with *ido* at the chromosome 5 QTL, this finding raises the possibility of a functional connection between the two loci through kynurenine metabolism.

The kynurenine pathway is the major route of tryptophan catabolism initiated by two different enzymes sharing an oxygenase domain, IDO and TDO. The pathway generates biologically active metabolites including kynurenic acid, quinolinic acid, and ultimately NAD⁺ ^42^. In resistant worms, *ido* expression was markedly reduced while *tdo* remained unchanged, suggesting that kynurenine metabolism is not globally suppressed but may be affected in specific cell populations where IDO is the dominant catabolic enzyme. Several independent lines of evidence support a role for this pathway in nematode drug response. Kynurenic acid antagonises glutamate binding in *H. contortus* membrane preparations^43^, metabolomic analyses identified reduced kynurenines in ivermectin-resistant *H. contortus*^44^, and profiling of the heartworm *Dirofilaria immitis* revealed differential kynurenine metabolite abundance between susceptible and resistant isolates^45^. Together, these observations suggest that kynurenine pathway dysregulation may represent a conserved adaptive response to macrocyclic lactone pressure across phylogenetically distinct nematodes. To investigate functional relevance, we tested whether kynurenic acid and quinolinic acid alter larval eprinomectin sensitivity. Kynurenic acid had no detectable effect, whereas quinolinic acid markedly potentiated eprinomectin activity (∼2.5-fold increase in sensitivity) in both susceptible and resistant larvae without affecting motility alone, confirming specificity of the effect. That both isolates were sensitised suggests quinolinic acid acts on the drug target pathway itself rather than specifically reversing resistance. The most parsimonious mechanistic explanation is indirect: quinolinic acid is a well-established regulator of glutamate homeostasis in vertebrates, increasing extracellular glutamate through stimulation of release and inhibition of uptake^46,47,48^. Since macrocyclic lactones act as allosteric activators of GluCl channels whose gating is functionally coupled to glutamate^49,50^, increased glutamate availability would be expected to enhance drug efficacy. Under this model, reduced IDO activity in resistant worms would lower quinolinic acid production, attenuate glutamatergic tone, and ultimately reduce GluCl activation by eprinomectin. Direct interaction of quinolinic acid with GluCls cannot be excluded – it shares physicochemical features with glutamate and has known ionotropic glutamate receptor activity – but this remains speculative. The absence of a kynurenic acid effect does not contradict this framework, as its pharmacokinetics in *H. contortus* are unknown and its regulatory effects on glutamatergic signalling may differ from those in vertebrates. Resolving these possibilities will require electrophysiological studies on recombinant *H. contortus* GluCls and direct measurement of glutamate dynamics in susceptible and resistant parasites.

Accumulating evidence suggests that macrocyclic lactone resistance more commonly involves broad neuronal rewiring and altered chloride homeostasis than selection at GluCl subunit genes^20,21^. Our findings are broadly consistent with this framework and extend it to eprinomectin, while revealing sex-specific and drug-specific dimensions not previously described. *cky-1*, a transcription factor proposed to regulate neuronal excitability and implicated in ivermectin resistance^21^, was expressed at low levels and modestly downregulated in resistant male and females. Nonetheless, its proposed downstream targets *pmk-1* and *pmk-2* were significantly underexpressed in resistant worms, suggesting residual *cky-1* regulatory activity at this locus. That the conserved chromosome 5 QTL may converge on partially distinct downstream effectors depending on drug, population, or resistance background is an important consideration for cross-study comparisons. *quiver* (HCON_00151600) and *sleepless* (HCON_00154990), neuroregulatory genes involved in cholinergic signalling and neuronal excitability, were significantly downregulated in resistant females. Broader transcriptomic patterns, including differential expression of voltage-gated ion channels, neurotransmitter transporters, and ligand-gated receptors, together with enrichment of locomotion-related phenotype ontologies, support a systems-level recalibration of neuronal circuitry under eprinomectin selection, extending prior findings^20^ to show that such reorganisation can be sex-specific and genetically coupled. *sulp-8* (HCON_00155080), orthologous to *C. elegans sulp-8* and related to the mammalian SLC26 anion transporter family, was one of only three genes consistently dysregulated across all three macrocyclic lactone resistance backgrounds, located within the chromosome 5 QTL with elevated variant allele frequencies. In mammals, disrupted SLC26 function can shift GABAergic signalling from inhibitory to excitatory and contribute to drug-resistant epilepsy^51,52^, suggesting that reduced *sulp-8* expression could similarly alter chloride gradients and modify GluCl-mediated currents in resistant nematodes – though functional characterisation will be required to confirm this. Finally, PGP-mediated drug efflux likely constitutes a complementary resistance layer: *Pgp-9.1* and *Pgp-9.2* were markedly overexpressed in resistant females, and *Pgp-9.1* in resistant males, consistent with their repeated identification across resistant nematode species^20,25^ and recent functional evidence for their role in macrocyclic lactone tolerance^53–55^. Rather than an alternative to the chromosome 4 and 5 signals, PGP overexpression likely acts in concert with neuronal remodelling, chloride dysregulation, and kynurenine pathway suppression – consistent with macrocyclic lactone resistance being shaped by the interaction of multiple physiological processes affecting both drug disposition and neuronal sensitivity.

In summary, this study provides the first integrated genomic and transcriptomic investigation of eprinomectin resistance in *H. contortus*. We confirmed and refined a major resistance locus on chromosome 5 shared across macrocyclic lactones, and identified a second QTL on chromosome 4 uniquely associated with eprinomectin-resistant populations and female-biased transcriptional enrichment. Multiple independent lines of evidence – genomic signatures of selection, recurrent transcriptional changes across resistant populations, and potentiation of eprinomectin activity by quinolinic acid – converge in implicating the kynurenine pathway in drug responsiveness. Establishing the causal roles of kynurenine metabolism, chloride homeostasis, and neuronal signalling, and whether these processes can improve anthelmintic efficacy or inform resistance diagnostics, remains an important goal for future work. Functional interpretation remains constrained by the limited functional annotation available for *H. contortus*, and the majority of genes identified under selection lack characterised homologues from which mechanistic hypotheses can be drawn. Nevertheless, our findings highlight key candidate genes and pathways, and provide a roadmap for future investigations into the genetic basis of anthelmintic resistance.

## Methods

### Ethics statement

All procedures for infection, maintenance, and euthanasia of animals associated with *Haemonchus contortus* sample collection were examined and approved by the Plateforme d’Infectiologie Expérimentale (PFIE), INRAE Centre Val de Loire, Nouzilly, France. All experiments were conducted in strict accordance with national ethics guidelines and approved by the local ethics committee for animal experimentation (Comité d’Ethique en Expérimentation Animale Val de Loire, CEEA VdL N°19) under protocol number APAFIS#17560. Where required, larval amplification was performed on experimentally infected ewes and rams under controlled conditions at the Toulouse National Veterinary School (UMR INRA/ENVT 1225, Interactions Hôtes-Agents Pathogènes, E3155527), under approval from the ethics committee for animal experimentation (APAFIS#39739).

### Haemonchus contortus isolates

The *H. contortus* isolates used in this study were originally collected and partially phenotypically characterised by Jouffroy *et al.*^5^ and Petermann *et al.*^7^. Isolates were obtained in 2020–2021 from five dairy sheep farms in a mountainous area of Southwest France. Flock management differed among farms: MOU used a collective summer pasture shared with other flocks, ARA used a non-collective pasture, and the remaining farms did not use summer pasture or mix flocks.

To obtain pure *H. contortus* populations, faecal samples containing parasite eggs were collected and coprocultures established to obtain infective L3-stage larvae. A drug selection step was then performed by experimentally infecting two 3- to 4-month-old lambs per isolate with L3. Animals harbouring eprinomectin-resistant isolates were treated subcutaneously with eprinomectin (0.2 mg/kg) 21 days post-infection, selecting against any residual susceptible worms and enriching for resistant genotypes. Susceptible isolates were maintained without drug treatment. Adult worms were subsequently recovered from the abomasum of each animal^7^, enabling collection of both male and female worms for RNA-seq, while L3 from coprocultures were used for Pool-seq.

### Phenotypic evaluation of the anthelmintic efficacy

The phenotypic drug efficacy of the three macrocyclic lactones (eprinomectin, ivermectin, and moxidectin) and levamisole was previously assessed in the laboratory^7^. Thiabendazole susceptibility was assessed using an egg hatch inhibition assay. Parasite eggs were recovered from fresh faecal samples by homogenisation in tap water followed by sieving, centrifugation, and flotation in saturated sodium chloride solution, as described by Coles *et al.*^56^. One hundred eggs in 79.2 µL were dispensed into each well of a 96-well plate alongside either 0.8 µL of thiabendazole solution or 0.8 µL of control solution (1% DMSO; total volume 80 µL). Final thiabendazole concentrations ranged from 5 ng/mL to 10 µg/mL, with each concentration tested in at least triplicate. Plates were sealed and incubated for 48 h at 23°C, then terminated by addition of 5 µL of Gram’s iodine solution (1 g iodine and 2 g potassium iodide in 100 mL distilled water) per well^57^. Unhatched eggs and released larvae were counted, and egg hatch inhibition was calculated as:

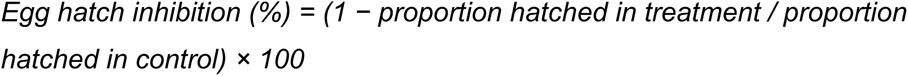

### Larval motility assay

Larval motility was assessed using the Worm MicroTracker One (WMiT; PhylumTech, Santa Fe, Argentina). Prior to assay, *H. contortus* L3 cuticles were removed by incubation in tap water supplemented with 0.15% NaOCl at 37°C for 15 min with agitation and vortexing every 5 min. Larvae were then washed four times with PBS (5 mL, 500 g, 5 min) to remove residual bleach and resuspended in LB medium. Eighty L3s per well were dispensed into 96-well flat-bottom plates in a final volume of 200 µL LB medium. Eprinomectin (0.01–100 µM) or 0.2% DMSO (negative control) was added, and plates were incubated at 37°C for 48 h in a humidified atmosphere (5% CO₂, ≥90% humidity) with or without the tested compounds kynurenic acid or quinolinic acid at 1 mM. Following incubation, larvae were exposed to light at room temperature for at least 30 min to restore motility before recording.

Motility was measured over a 15 min period by dual infrared microbeam interruption counts, then normalised to the mean of control wells to derive percentage motility inhibition. All experiments were performed in triplicate.

### DNA extraction, library preparation and sequencing

DNA was isolated from pools of 200 L3 larvae. Frozen pellets were resuspended in 300 μL lysis buffer (200 mM NaCl, 100 mM Tris-HCl, 30 mM EDTA pH 8, 0.5% SDS) with 20 μL proteinase K (20 mg/mL) and incubated at 55°C for 2 h. RNA was removed by addition of 10 μL RNase A (10 mg/mL) and incubation at 37°C for 10 min. DNA was purified by phenol/chloroform/isoamyl alcohol (25:24:1) extraction, precipitated overnight at −80°C with 0.1× volume sodium acetate (pH 5.5), 3× volume 100% ethanol, and 2 μL glycogen, then pelleted by centrifugation (14,000 g, 4°C, 5 min). The pellet was washed with 500 μL 70% ethanol, air-dried, and resuspended in ≥10 μL EB buffer. Genomic DNA was stored at 4°C. Libraries were prepared using the NEBNext Ultra II DNA Library Prep Kit for Illumina with dual indexing, pooled as a 6-plex, and sequenced on a single Illumina NovaSeq 6000 XP lane using 150 bp paired-end chemistry.

### RNA extraction, library preparation for RNA-seq

Adult worms from one donor each for the five *H. contortus* isolate populations were used for RNA-seq. Total RNA was isolated from three batches of 20 male or female worms per donor using a standard TRIzol (Thermo Fisher Scientific) extraction protocol. Each RNA sample underwent DNaseI (Qiagen) treatment in solution, followed by purification on a RNeasy Mini Spin Column (Qiagen), before freezing at - 80°C. Total RNA (1 μg) from each sample was used for NEBNext PolyA selection and Ultra Directional RNA Library preparation (Illumina). Oligo-dT pulldown RNA sequencing libraries were prepared using a NEBNext Ultra II Directional RNA Library Prep Kit for Illumina. Individually barcoded libraries were pooled into a 36-plex and sequenced on two lanes of an Illumina NovaSeq 6000 SP using 100-bp paired-end chemistry. Sequencing data for both DNA and RNA, including sample accession numbers and sequencing lane IDs, are provided in Supplementary Data 1.

### Genomic analyses

#### Read mapping

Raw sequence data were assessed using FastQC (https://www.bioinformatics.babraham.ac.uk/projects/fastqc/) and summarised using MultiQC v.1.17^58^. We used a Nextflow mapping pipeline (mapping-helminth/v1.0.8; https://github.com/sanger-pathogens/mapping-helminth) to map raw sequencing reads to the *H. contortus* genome (WormBase ParaSite release 18 PRJEB506;^59^). Briefly, GATK v.4.1.4.1^60^ was used to convert paired-end reads to a interleaved ubam using FastqToSam and processed using MarkIlluminaAdapters, before mapping to the *H. contortus* genome using minimap2 v.2.16^61^. SAMtools^62^ and Sambamba^63^ were used to sort and mark duplicates in the resulting BAM output. MultiQC was again used to summarise mapping statistics generated using SAMtools flagstats.

#### Analyses of genetic variation

We analysed genetic variation to (i) detect patterns of variation change throughout the genome via pairwise comparisons of isolates, and (ii) identify single-nucleotide changes in genes that are associated with resistance within each isolate. To do this, we started with BAM files. To compare genetic differentiation between isolates, we used grenedalf v.0.6.3^64^ to measure genetic differentiation (*F*_ST_) in 5 kb windows with a step size of 2.5 kb. We compared susceptible-vs-resistant, susceptible-vs-susceptible, and resistant-vs-resistant groups, keeping consistent parameters to enable comparison (--method unbiased-nei --sam-min-map-qual 30 --sam-min-base-qual 30 --filter-sample-min-count 2 --filter-sample-min-read-depth 30 --filter-sample-max-read-depth 300 --pool-sizes 1000 --window-type interval --window-average-policy valid-loci). For zoomed-in comparisons of peaks of differentiation, we used a rolling mean *F*_ST_ in 200 kb windows.

Per-sample variant calling was performed from BAM files filtered for mapping quality >20 using bcftools v1.23 (mpileup, then call, then norm -m for biallelic normalisation^65^). Variant effects were predicted using SnpEff v5.1^66^. Annotated VCFs were merged into a multi-sample variant table, with sample-level genotype fields converted to alternative allele frequencies (AF = AD_alt/DP); missing, zero-depth, or non-interpretable calls were coded as 0. Variants were filtered by allele-frequency contrast between phenotypic groups: those with AF > 0.1 in any susceptible sample were removed. Because ivermectin resistance has previously been described as dominantly inherited^22,67^, heterozygous individuals may be resistant; variants were therefore retained if AF > 0.4 in at least one resistant sample. This threshold was intended to capture variants that may remain at intermediate population frequencies while being enriched by positive selection in resistant populations. The resulting dataset represents variants rare or absent in susceptible samples and present at moderate-to-high frequency in resistant samples.

Filtered variants were intersected with gene and exon intervals from the GTF file, retaining only those overlapping annotated features. For each gene, variant counts were calculated, density was normalised as variants per kilobase, and allele frequencies were averaged across all assigned variants. Mean AFs were calculated separately for resistant and susceptible samples, and the plotted statistic was μAFR − μAFS, such that each point represents one gene, with higher values indicating variants enriched in resistant isolates. This gene-level summary facilitates screening for candidate resistance loci across the genome.

### Transcriptomic analyses

#### Raw read processing, read mapping, and gene counts

As with genomic data, the transcriptomic data were first assessed using FastQC and MultiQC. Raw FastQ files were trimmed for adapter contamination using Cutadapt v.4.3^68^ with default settings. Reads were aligned to the *H. contortus* reference genome (WormBase ParaSite release 18 PRJEB506;^59^) using Hisat2 v.2.2.1^69^. The resulting SAM files were converted to BAM and indexed using SAMtools v1.19. Duplicates were marked with Sambamba v.1.0.1^63^. For mapping QC, we used samtools flagstat and stats.

RNA-seq data from a previously published study^19^ on ivermectin and moxidectin transcriptomic responses were used to compare against the transcriptomes of eprinomectin-sensitive and resistant isolates generated here. For consistency, raw data from both previous studies were processed as described above.

The gene counts table was generated with featureCounts from the Rsubread v.2.16.1^70^ library of Bioconductor in R. A normalised counts table was generated in MRM with DESeq2 v.1.44.0^71^ from Bioconductor in R.

#### Sample concordance

Principal component analysis was performed on normalised, log-transformed read counts using the R base function prcomp, and visualised using the plotly library^72^. Sample clustering was assessed using pheatmap (https://github.com/raivokolde/pheatmap).

One male sample, BUN-M3, was excluded prior to differential expression analysis on the basis of poor concordance with other male samples (Supplementary Fig. 10).

Poisson distance analysis confirmed that BUN-M3 was markedly more dissimilar from other males than any other pairwise comparison. Examination of the most variable genes revealed elevated expression of three probable vitellogenin orthologs (HCON_00099370, HCON_00184500, HCON_00188550) – yolk protein genes with otherwise negligible expression in male samples – approaching levels observed in female worms. This pattern is consistent with female material contamination of this sample. BUN-M3 was therefore excluded from all downstream analyses.

#### Differential expression

Differential gene expression analyses were performed on raw counts for males and females separately, stratified by resistance status of *H. contortus* isolates, using DESeq2 (v.1.46.0). Differentially expressed genes (DEGs) were defined as those with an adjusted *p*-value < 0.05 and Log_2_ fold change (Log_2_FC) > 0.58 or <-0.58 (1.5-fold change). From the raw DGE table, interactive volcano plots were produced using Plotly (as mentioned above).

#### Gene-set enrichment

Gene-set enrichment analyses were performed on DEGs using all genes with a non-zero total read count as a background based on Gene Ontology database. The analyses were carried out using gprofiler2 (biit.cs.ut.ee/gprofiler), with default parameters and *H. contortus* as the set organism. Gene Ontology and Phenotype Enrichment Analysis with *C. elegans* orthologs were performed using the gene-set and phenotype Enrichment Analysis tools from Wormbase (https://wormbase.org)^73^ with a threshold Q-value of 0.1. The output files were processed, and data visualized using ggplot2.

#### Real-time quantitative PCR (RT-qPCR)

Total RNA from each isolate (1 µg), drawn from the same extractions used for RNA-seq, was treated with ezDNase Enzyme (Thermo Fisher Scientific) to remove genomic DNA, then converted to cDNA using the SuperScript IV VILO Master Mix (Thermo Fisher Scientific) following the manufacturer’s instructions. qPCR was performed using PowerTrack SYBR Green Master Mix (Thermo Fisher Scientific), with primers targeting *cky-1* (HCON_00155390; Forward: 5′-CCGAGACCAGATCAATGTCG-3′, Reverse: 5′-CACACTGTCTTCGGCTATCG-3′) and the reference gene *β-actin* (HCON_00135080; Forward: 5′-CCAGTTGGTGACGATTCC-3′, Reverse: 5′-GGGTTTGCTGGAGATGACG-3′).

Reactions were performed in triplicate on a QuantStudio™ 6 Pro (Thermo Fisher Scientific) using the following cycling conditions: 95°C for 2 min, followed by 40 cycles of 95°C for 20 s, 55°C for 20 s, and 60°C for 20 s. Melt curve analysis (95°C for 15 s, 60°C for 1 min, ramping to 95°C at 0.075°C/s) confirmed a single amplification product per primer pair. gDNA digestion, no-RT, and no-template controls were included in each run, confirming amplification specificity. Relative expression was calculated using the ΔΔCt method with QuantStudio™ Design & Analysis Software v2.8.0.

## Data availability

Sequencing data generated in this study have been deposited in the European Nucleotide Archive under the study accessions ERP144506 (whole genome data) and ERP144509 (transcriptomics data), and are described in full in Supplementary Data 1.

## Code availability

The code used in the study is available via GitHub at https://github.com/stephenrdoyle/hcontortus_eprinomectin and is archived under a stable DOI at Zenodo.

## Acknowledgements

The authors would like to thank the “Plate-forme d’Infectiologie Expérimentale (PFIE)” that contributed to amplify *H. contortus* isolates, and the Parasites and Microbes Operations and Informatics teams at the Wellcome Sanger Institute for logistical and computational support.

This work was funded by Carnot France Future Elevage (2025) https://www.francefuturelevage.com/ (ANTHERIN, GENOPAR), the European ICRAD program (HARTEMIS ANR-24-ICRD-0003-01), the Wellcome Trust (UK) through core funding to the Wellcome Sanger Institute (UK) (220540/Z/20/A), and a UKRI Future Leaders Fellowship (MR/T020733/1). For the purpose of Open Access, the authors have applied a CC BY public copyright Licence to any Author Accepted Manuscript version arising from this submission.

## Author contributions

A.L. and S.R.D. designed the study and analysed data. J.P. and P.J. oversaw worm sample collection. J.P. and F.G performed the drug efficacy phenotyping. E.C. and G.S. extracted DNA; R.B. extracted RNA. S.R.D. performed RT-qPCR experiments. R.B. and E.G. performed *C. elegans* experiments. R.B., R.L., and S.R.D. performed bioinformatic analyses, analysed data, and generated figures. R.B, R.L, S.R.D, and A.L drafted the manuscript. All authors have read and approved the final version of the manuscript.

