## Supplementary Information for "Sex-biased transcriptional rewiring of neuronal circuits associated with eprinomectin resistance in *Haemonchus contortus*"

**Supplementary Table 1. Anthelmintic resistance profiles of *Haemonchus contortus* isolates.**

| | Eprinomectin (µM) | | | | Ivermectin (µM) | | | | Moxidectin (µM) | | | | Thiabendazole (µM) | | | |
|---|---|---|---|---|---|---|---|---|---|---|---|---|---|---|---|---|
| Isolate | $IC_{50}$ | SD | RF | *n* | $IC_{50}$ | SD | RF | *n* | $IC_{50}$ | SD | RF | *n* | $IC_{50}$ | SD | RF | *n* |
| CHI | 0.33 | 0.21 | 1.0 | 4 | 0.37 | 0.25 | 1.0 | 4 | 0.11 | 0.06 | 1.0 | 4 | 0.28 | 0.01 | 1.0 | 1 |
| LUC | 0.41 | 0.23 | 1.0 | 4 | 0.18 | 0.15 | 1.0 | 4 | 0.09 | 0.06 | 1.0 | 4 | 0.22 | 0.01 | 1.0 | 1 |
| ARA | 18.81 | 15.07 | 63.7 | 2 | 3.71 | 2.25 | 13.5 | 2 | 0.34 | 0.19 | 4,2 | 2 | 4.64 | 0.20 | 17.8 | 1 |
| BUN | 40.40 | 29.41 | 136.9 | 3 | 0.77 | 0.15 | 2.8 | 3 | 0.24 | 0.10 | 3.0 | 3 | 0.14 | 0.10 | 0.5 | 1 |
| MOU | 25.82 | 11.51 | 87.5 | 2 | 2.05 | 0.73 | 7.4 | 2 | 0.32 | 0.21 | 4.0 | 2 | 6.31 | 0.21 | 25.2 | 1 |

For macrocyclic lactones, $IC_{50}$ were determined using worm microtracker motility assays. Values are the mean and standard deviation (SD) of 2-4 experiments. For thiabendazole, $IC_{50}$ were determined from egg hatch assays. Values are the mean of 6 replicates for each concentration point (one experiment). Resistance Factors (RF) were calculated as follows: $IC_{50}$(isolate) / ([$IC_{50}$(CHI) + $IC_{50}$(LUC)] / 2).

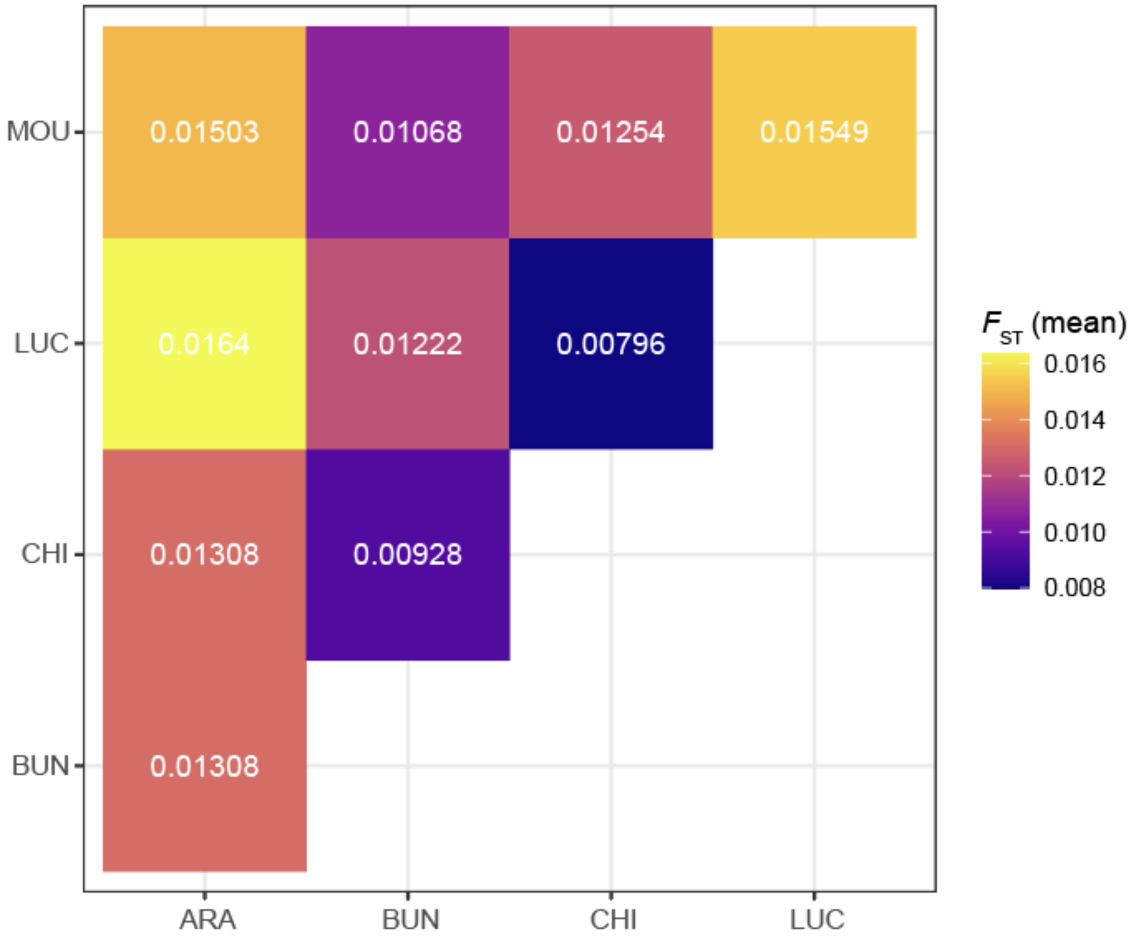

**Supplementary Fig. 1: Pairwise genetic differentiation ($F_{ST}$) between all populations.**
Upper triangular heatmap of genome-wide mean pairwise $F_{ST}$, computed from windowed $F_{ST}$ values, for all pairs of isolates: eprinomectin-susceptible LUC and CHI, and eprinomectin-resistant ARA, BUN, and MOU. The lowest differentiation is observed between the susceptible populations CHI and LUC, whereas LUC and the resistant isolate ARA show the greatest divergence, consistent with their greater geographic separation.

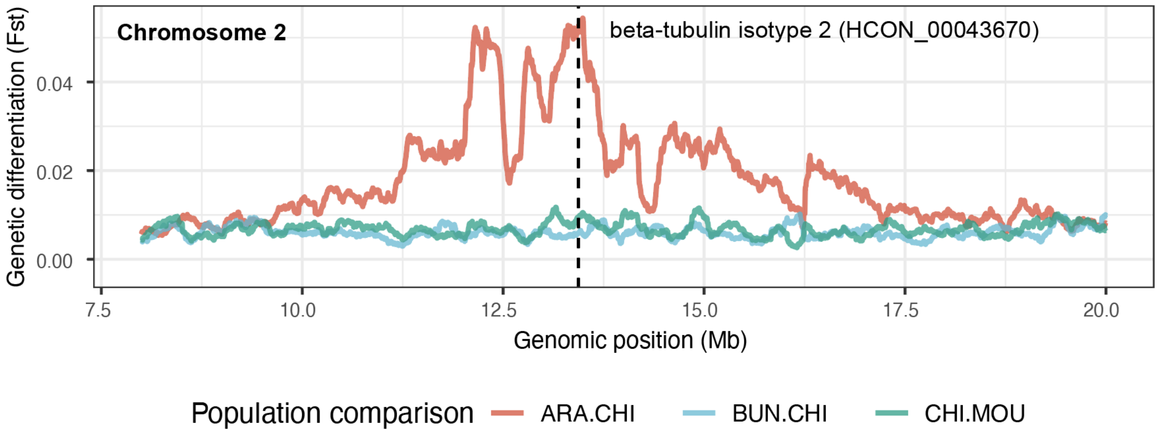

**Supplementary Fig. 2: A peak of differentiation on chromosome 2 at the beta-tubulin isotype 2 locus is observed only in the ARA population.**
Smoothed windowed $F_{ST}$ (5,000-bp windows, 2,500-bp overlap, smoothed over 200-kb sliding windows) across a region of chromosome 2 for three pairwise comparisons between each resistant isolate (ARA, BUN, MOU) and the susceptible CHI population. The vertical dashed line indicates the position of the beta-tubulin isotype 2 gene, which has previously been associated with benzimidazole resistance. An elevated peak of differentiation in ARA–CHI suggests selection at this locus in ARA; however, no known resistance-associated variants were identified within the gene.

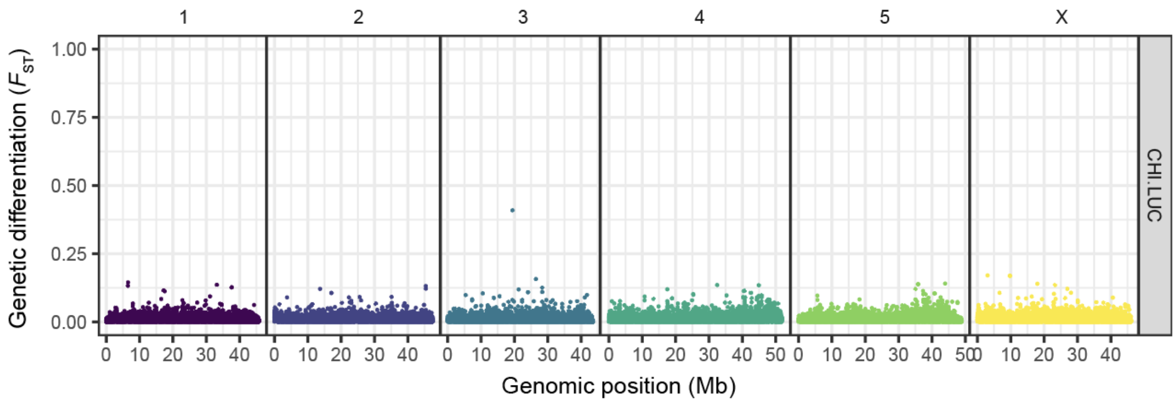

**Supplementary Fig. 3: Genome-wide $F_{ST}$ between eprinomectin-susceptible populations reveals low background differentiation.**

Pairwise genetic differentiation ($F_{ST}$ between eprinomectin-susceptible CHI and LUC populations across all chromosomes. Each point represents a 5,000-bp window (2,500-bp overlap), with $F_{ST}$ calculated as the mean across all variants within that window. Consistent with the low genome-wide mean $F_{ST}$ shown in Supplementary Fig. 1, differentiation between these populations is uniformly low across the genome. The y-axis is scaled to 1.0 to enable direct comparison with figures showing resistant population comparisons.

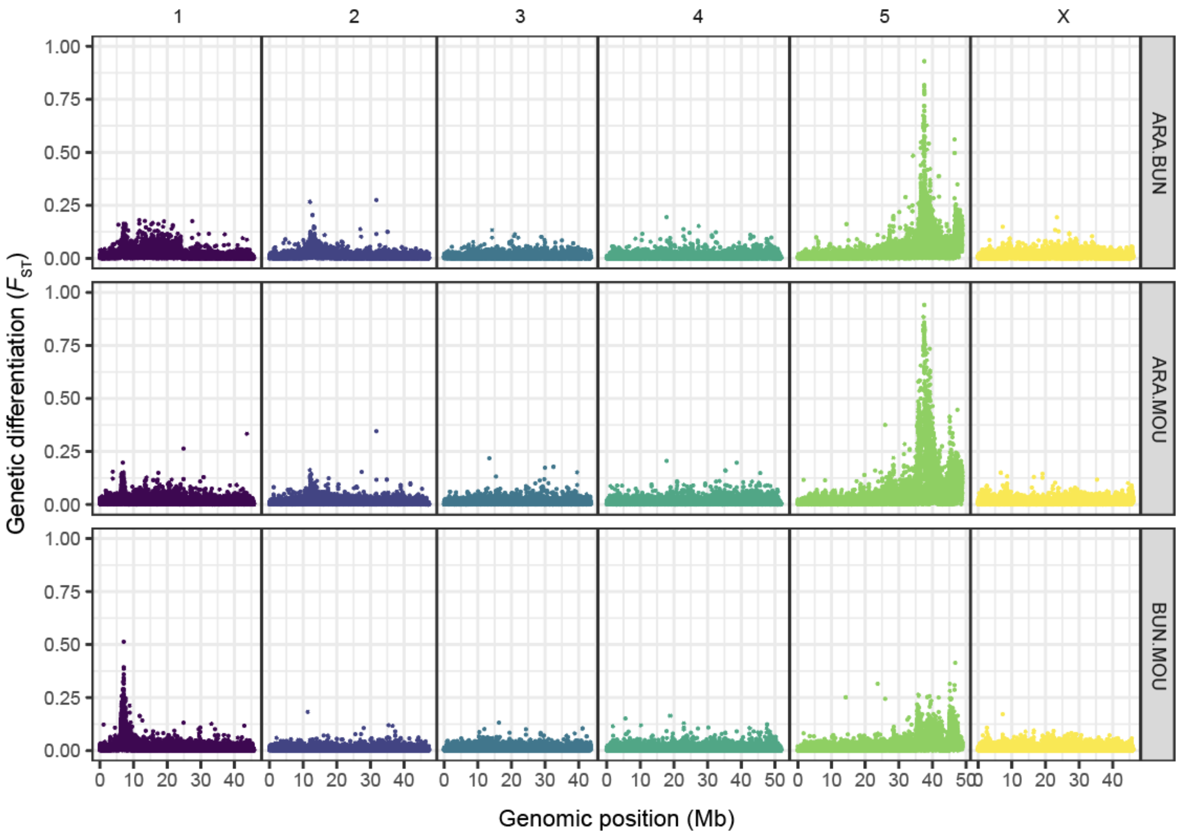

**Supplementary Fig. 4: Pairwise $F_{ST}$ between eprinomectin-resistant populations reveals a shared peak of differentiation at the end of chromosome 5.**

Genome-wide pairwise genetic differentiation ($F_{ST}$) between eprinomectin-resistant populations ARA, BUN, and MOU, shown for all three pairwise combinations (rows, top to bottom: ARA–BUN, ARA–MOU, BUN–MOU). Each point represents a 5,000-bp window (2,500-bp overlap), with $F_{ST}$ calculated as the mean across all variants within that window. A pronounced peak of differentiation is visible at the distal end of chromosome 5 in all three comparisons, reaching $F_{ST}$ values approaching 1.0 in ARA–BUN and ARA–MOU, suggesting strong selective differentiation at this locus among resistant populations.

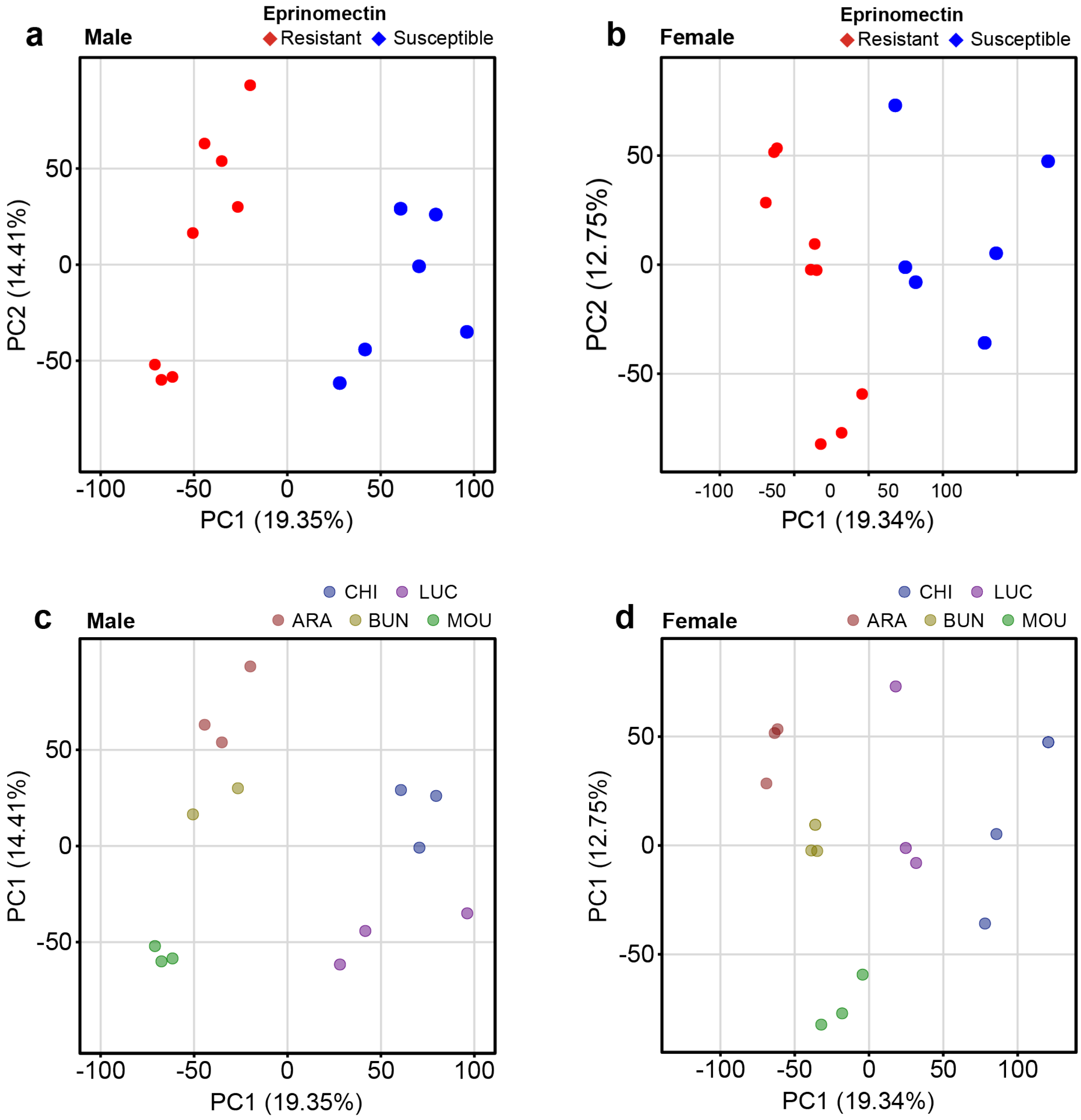

**Supplementary Fig. 5: PCA of transcriptomic data separates eprinomectin-resistant and susceptible adult *Haemonchus contortus* in both sexes.** Principal component analysis (PCA) of regularised log-transformed expression data from 14,112 genes, shown separately for **a**. and **c.** males and **b**. and **d.** females. Each point represents an individual worm sample; resistant (R) populations are shown in red and susceptible (S) in blue in **a.** and **b.**, while samples are colored by farm location in **c**. and **d.** PC1 separates resistant from susceptible populations in both sexes, accounting for 19.35% and 19.34% of variance in males and females, respectively.

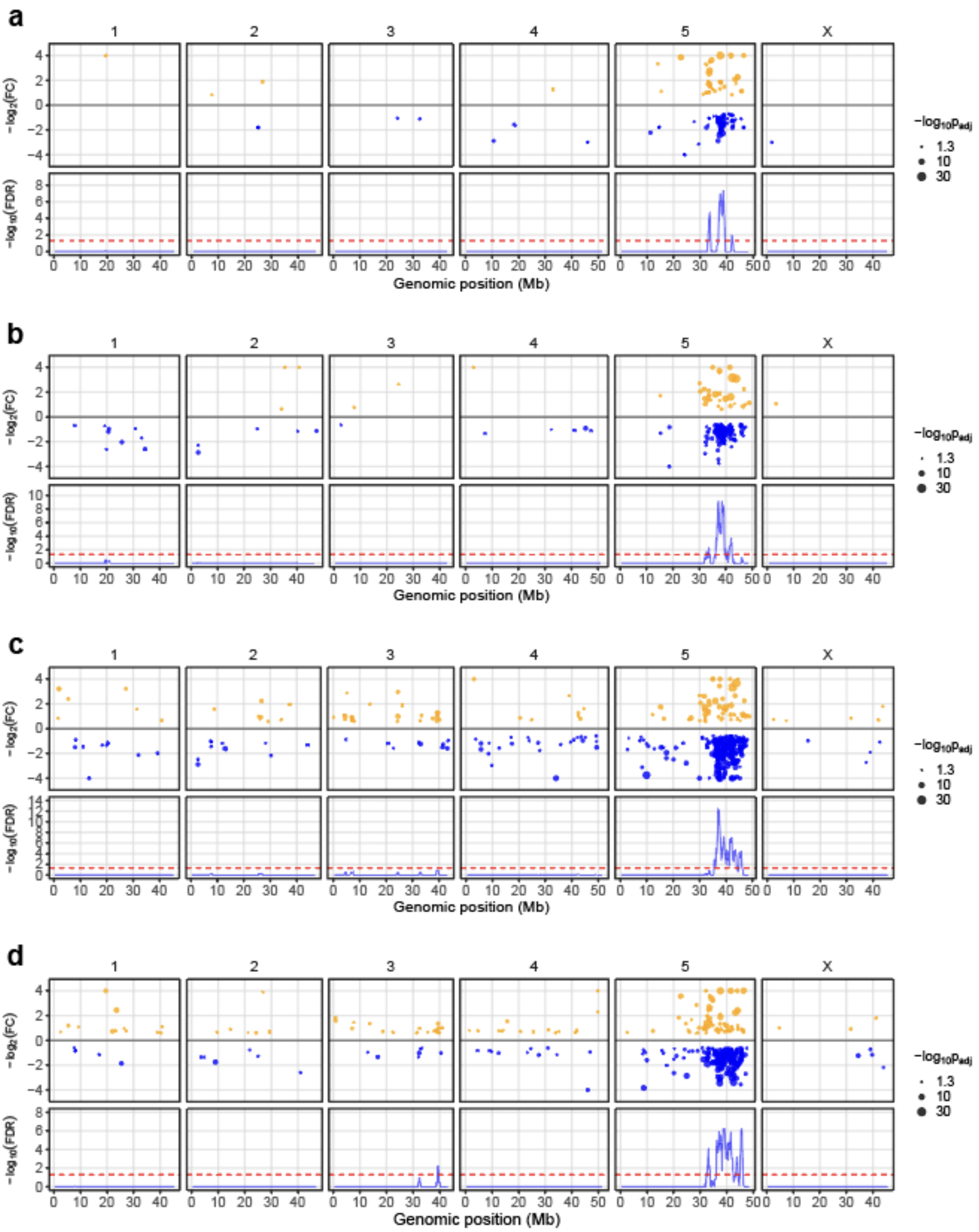

**Supplementary Fig. 6: Reanalysis of ivermectin and moxidectin transcriptomic data reveals consistent enrichment of differentially expressed genes within regions of genomic differentiation on chromosome 5.**
Genome-wide karyoplots showing the genomic distribution of differentially expressed genes (DEGs) in Ivermectin-selected resistant males (**a**) and females (**b**), and Moxidectin-selected resistant males (**c**) and females (**d**), obtained from a reanalysis of previously published transcriptomic datasets[1]. Yellow and blue points indicate genes significantly upregulated or downregulated, respectively, in resistant isolates

relative to susceptible isolates. Each point represents a DEG (adjusted p-value < 0.05), and point size reflects the level of statistical significance. For visualisation purposes, genes with log2 fold-change values outside the [-4, 4] range were truncated to the corresponding boundary values. Blue curves show the significance of hypergeometric enrichment tests assessing the clustering of DEGs within 1 Mb sliding genomic windows with a 100 kb step size. False discovery rates (FDRs) were calculated using the Benjamini–Hochberg correction, and the red dotted lines indicate the significance threshold corresponding to FDR = 0.05.

**Exploring *cky-1* expression in susceptible and eprinomectin-resistant isolates**

A leading candidate gene, *cky-1*, associated with drug resistance to the macrocyclic lactone ivermectin, was recently identified in genomic and transcriptomic analyses of genetic crosses between susceptible and ivermectin-resistant isolates of *H. contortus*[2,3]. Support for this came from several lines of evidence:

1. Strong evidence of selection around the gene as a QTL centred approximately 37.5 Mb on chromosome 5.
2. Significant increase in gene expression of *cky-1* in ivermectin-resistant *H. contortus* isolates relative to susceptible isolates by RNA-seq and RT-qPCR in genetic crosses and other susceptible and resistant isolates.
3. Significant increase in gene expression of *cky-1* in ivermectin-resistant *Teladorsagia circumcincta* isolates relative to susceptible isolates.
4. RNAi knockdown and balanced deletion of *cky-1* in *Caenorhabditis elegans* led to hypersensitivity to ivermectin.

Considering these lines of evidence, *cky-1* was a candidate of interest as a potential driver of eprinomectin resistance in this study. Genomic comparison of susceptible and eprinomectin-resistant isolates in Fig. 2 shows strong genetic differences, represented by a peak of differentiation, that overlap significantly with *cky-1*, consistent with previous work on ivermectin-resistant isolates. However, our RNA-seq transcriptomic analyses failed to detect any reads associated with *cky-1*. This is not particularly surprising, given that *cky-1* is a transcription factor with very low expression, is tightly regulated, and is expressed in only a few cells in the organism (inferred from *C. elegans*).

To explore this further, we analysed *cky-1* expression by RT-qPCR using the same RNA samples used for RNA-seq. In these experiments, we did detect low levels of *cky-1* expression in all samples relative to the internal *β-actin* control (Supplementary Fig. 7a). Almost all comparisons between susceptible and resistant populations in both sexes were non-significant, suggesting no difference in *cky-1* expression based on treatment groups. Exceptions were seen in ARA females, with lower expression of *cky-1* relative to both CHI (*P*[adjusted] = 0.0107) and LUC (*P*[adjusted] = 0.0268), and ARA males with LUC (*P*[adjusted] = 0.00346). Similarly, BUN males showed lower *cky-1* relative to LUC (*P*[adjusted] = 0.0111), but not CHI (*P*[adjusted] = 0.739). When all populations were considered together, a decrease in *cky-1* expression was observed in resistant females (*P*[adjusted] = 0.0138) and males (*P*[adjusted] = 0.0127) relative to susceptible females and males (Supplementary Fig. 7b). Collectively, these results do not support the observation seen in ivermectin-resistant *H. contortus*. Our data show, at most, modest and inconsistent differences in *cky-1* expression between resistant and susceptible isolates, with a tendency toward decreased rather than increased expression in resistant populations.

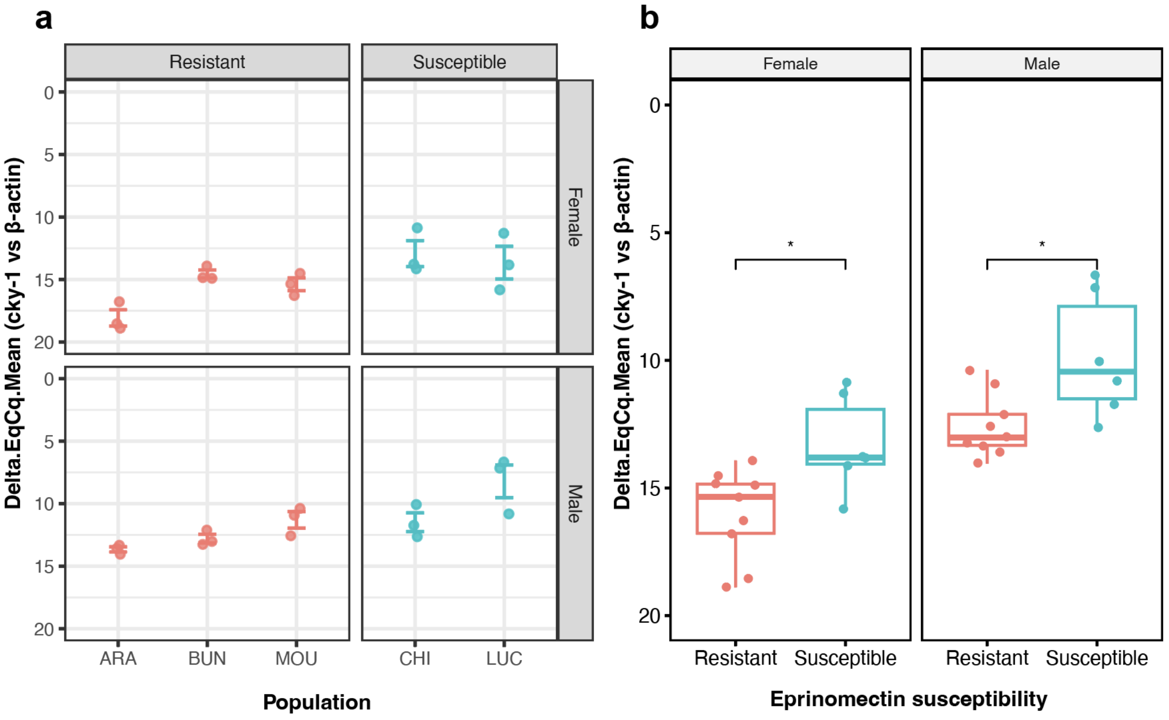

**Supplementary Fig. 7: RT-qPCR analysis of *cky-1* expression in susceptible and eprinomectin-resistant isolates.**
Relative expression of *cky-1* normalised to *β-actin* was measured by RT-qPCR. Each point represents the mean Delta.EqCq (difference in amplification cycles between *β-actin* and *cky-1*) for one biological replicate (n = 3 per population per sex), each measured in technical triplicate. Note that higher Delta.EqCq values indicate lower relative expression. **a.** Relative *cky-1* expression by population, sex, and treatment group; error bars show standard error of the mean. **b.** Relative *cky-1* expression pooled by treatment group and sex. Boxplots show the median, the interquartile range, and whiskers extending to 1.5× the IQR. Statistical comparisons used a t-test; * indicates $P < 0.05$; ns, not significant.

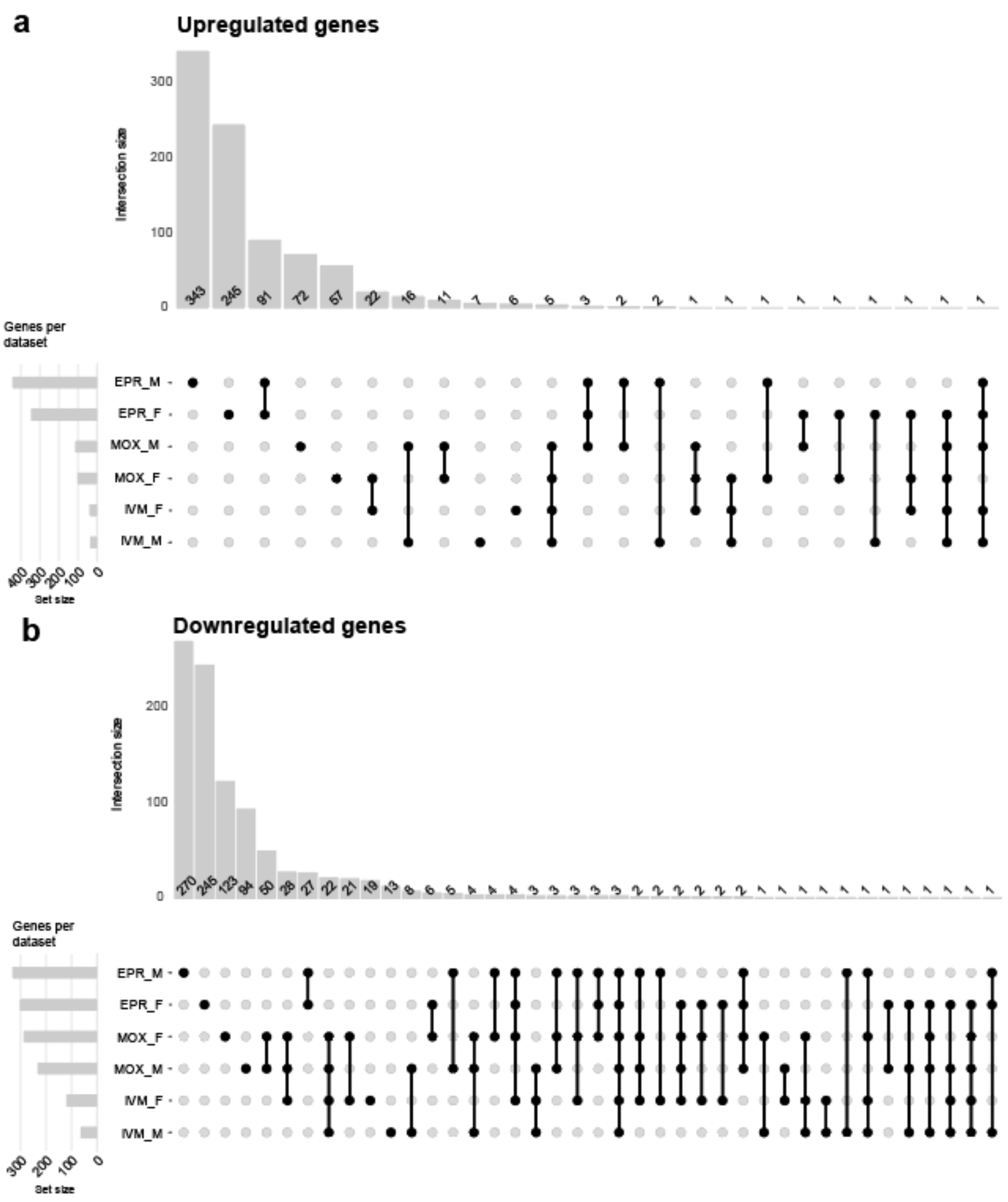

**Supplementary Fig. 8: Comparison of differentially expressed genes across macrocyclic lactone-resistant *Haemonchus contortus* isolates.**

UpSet plots showing the overlap of significantly upregulated (**a**) and downregulated (**b**) genes identified in sex-specific comparisons between susceptible and resistant isolates from the present eprinomectin (EPR) study and from the ivermectin (IVM)- and moxidectin (MOX)-selected isolates reported by McIntyre *et al.* [1]. Each set corresponds to the list of differentially expressed genes obtained for a given sex and drug selection regime. Intersections highlight genes showing shared patterns of differential expression across macrocyclic lactone-resistant isolates, whereas unique sets represent treatment- or sex-specific transcriptional responses.

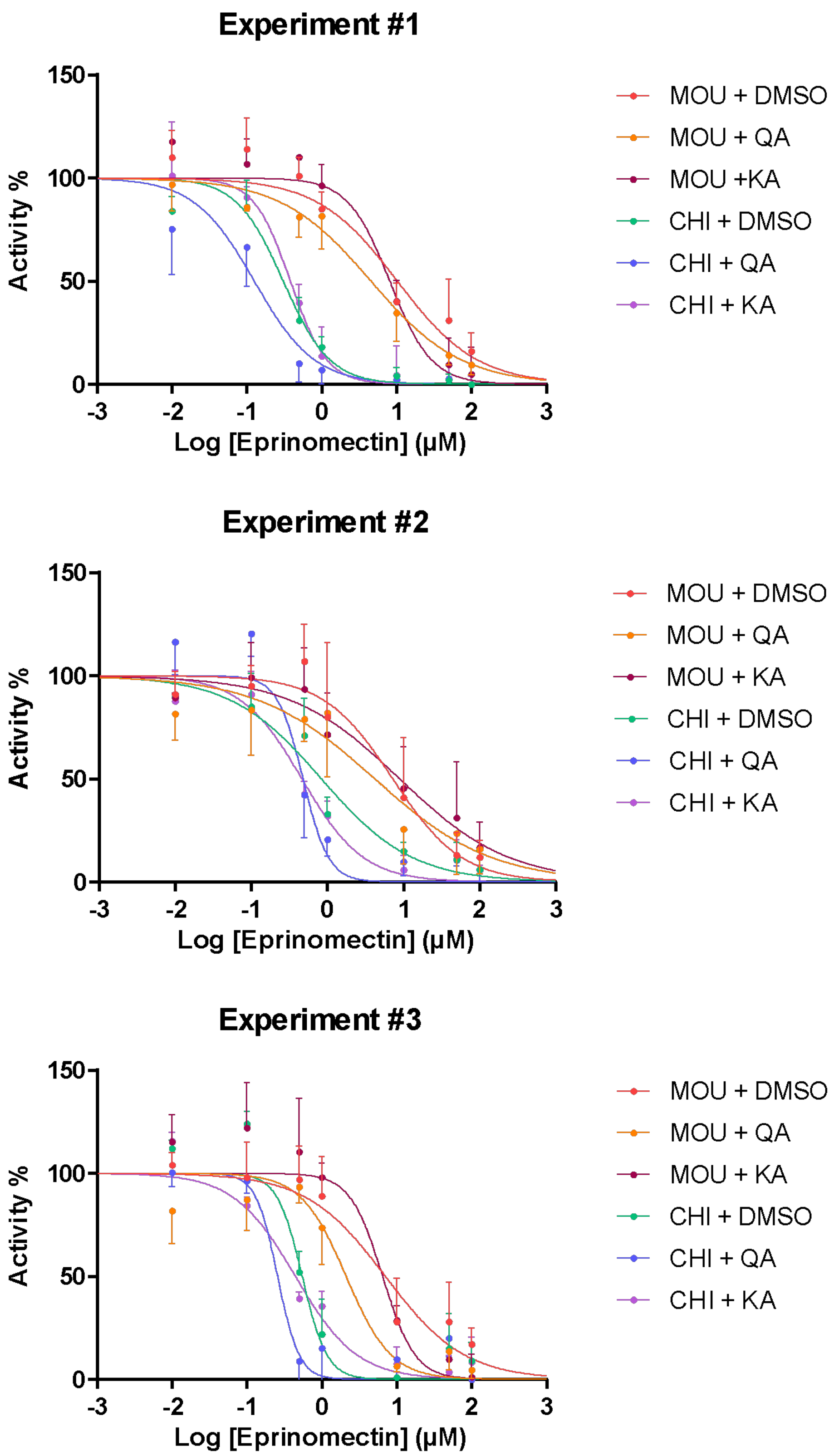

**Supplementary Fig. 9: Experimental replicates of motility assay in dose-response assays to eprinomectin with kynurenic acid (KA) or quinolinic acid (QA)**

Replicates correspond to the data shown in Fig. 6b.

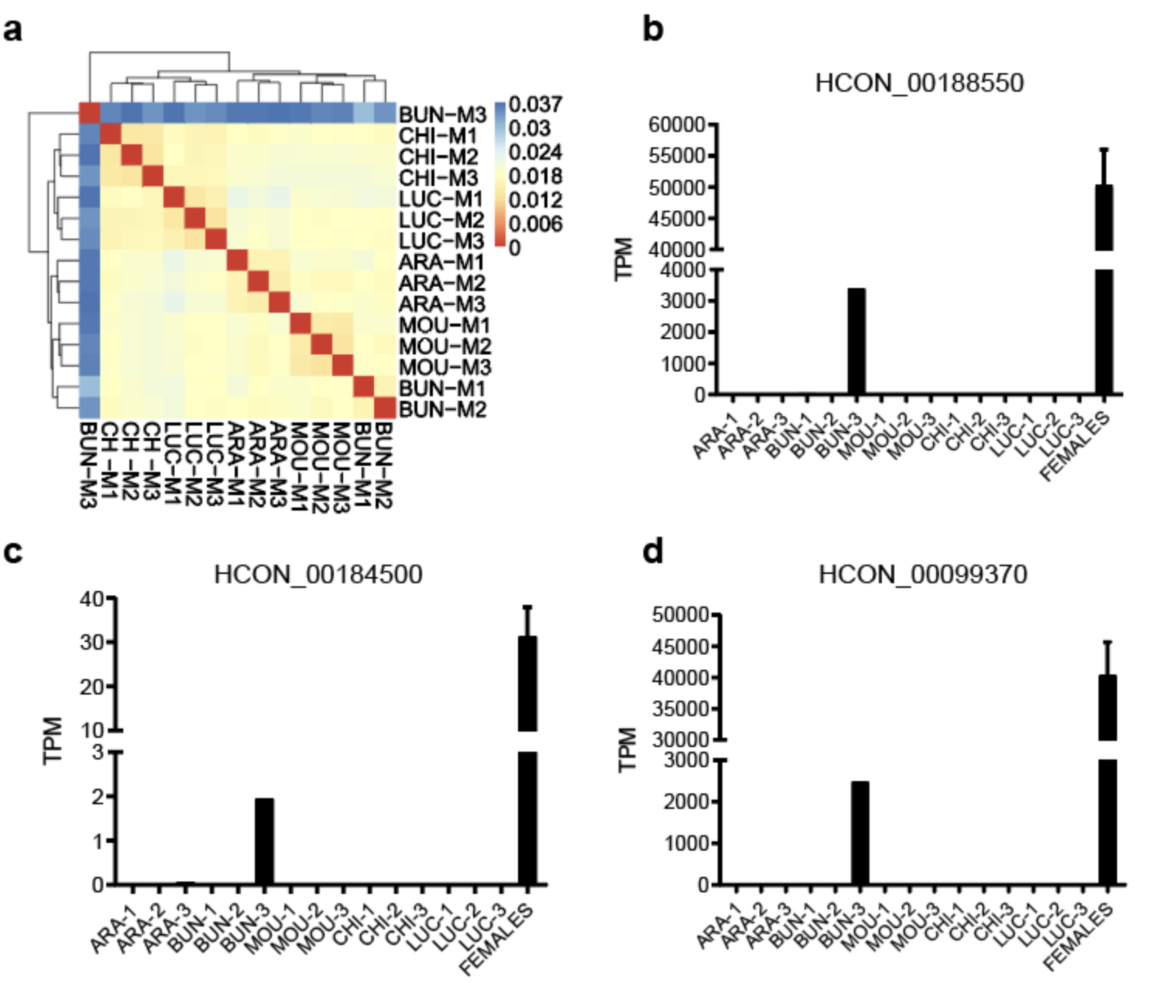

**Supplementary Fig. 10: Analyses of the male BUN-M3 sample reveal likely contamination with female-specific genes.**
**a**. Clustered heatmap of gene expression in male samples revealed that BUN-M3 was an outlier relative to all male samples. Colours represent the distance (1 - Pearson correlation) between pairwise samples, with red indicating similar samples and blue indicating distant samples. **b,c,d**, Analyses of three vitellogenin domain-containing protein genes that should show female-specific gene expression among all male samples reveal low levels of expression in BUN-M3, suggesting a mixed male and female gene expression profile. Expression levels in transcripts per million (TPM) of vitellogenin domain-containing protein genes (**b**) HCON_00188550, (**c**) HCON_00184500, (**d**) HCON_00099370.
